# Aromatase in adipose tissue exerts an osteoprotective function in male mice via phosphate regulation

**DOI:** 10.1101/2024.06.24.600344

**Authors:** Aoi Ikedo, Michiko Yamashita, Maiko Hoshino, Yosuke Okuno, Megumi Koike, Minori Uga, Kazuya Tanifuji, Hiroko Segawa, Seiji Fukumoto, Yuuki Imai

## Abstract

Aromatase contributes to maintenance of bone mass because male patients with loss-of-function mutations of *CYP19A1* exhibit bone loss. Treatment with aromatase inhibitor also causes bone loss in men and post-menopausal women, suggesting that part of the anabolic effect of testosterone in men is dependent on estradiol (E2) biosynthesized by aromatase in non-gonadal tissues. It remains unclear how locally biosynthesized E2 contributes to maintenance of bone mass. We examined the function of aromatase in local tissues rather than gonads using cell-type specific aromatase knockout (KO) mice. Because osteoblast-specific aromatase KO mice exhibited no bone phenotype, we focused on adipose tissue, known as a reservoir of steroid hormones and analyzed the bone phenotypes of adipose tissue-specific aromatase KO (*Aro*^Δ*aP2*^) mice. Sixteen-week-old male *Aro*^Δ*aP2*^ mice exhibited significantly lower bone mineral density in tibia and femur, especially in trabecular bone, than controls. Bone histomorphometry showed that *Aro*^Δ*aP2*^ mice exhibited an insufficient calcification bone phenotype with increased osteoid volume and width, and decreased osteoclast area and numbers. Moreover, serum phosphate, renal phosphate reabsorption and FGF23 were significantly lower in *Aro*^Δ*aP2*^, suggesting that the insufficient calcification phenotype in *Aro*^Δ*aP2*^ was not caused by excessive FGF23 activities. Finally, we analyzed NaPi2a and NaPi2c, phosphate transporters localized in the kidney, and found that protein levels in renal brush border membrane vesicles were lower in *Aro*^Δ*aP2*^. These results indicate that estrogens locally biosynthesized by aromatase in adipocytes can play a significant role in bone mass maintenance via regulation of phosphate reabsorption in the kidney by NaPi2.

**Impact Statement:** Male adipocyte-specific aromatase KO mice (*Aro*^Δ*aP2*^) exhibited bone loss and increased osteoid due to decreased serum phosphate and urinary phosphate reabsorption as a result of reduced NaPi2 proteins expression in the kidney.

## Introduction

Postmenopausal osteoporosis is a major medical problem that reduces the quality of life in elderly women (Rosen, 2005). Estrogen deficiencies can induce postmenopausal osteoporosis due to the lack of various physiological functions of estrogen/estrogen receptor signaling, such as maintenance of osteoclastic life span (Martin-Millan et al., 2010; Nakamura et al., 2007), maintenance of osteoblast/osteocyte functions by Wnt signaling (Almeida et al., 2013; Doolittle et al., 2022; Kondoh et al., 2014), inhibition of inflammatory cytokine production (Pacifici, 2008), and negative feedback of FSH secretion (Sun et al., 2006). Even in older men, low serum levels of estradiol (E2) (Mellström et al., 2008) as well as testosterone (Shahinian, Kuo, Freeman, & Goodwin, 2005) can lead to significant risks of fractures. Estrogens are synthesized from cholesterols by the activity of converting enzymes, including aromatase (*Cyp19a1*), mainly in the gonads. The *Cyp19a1* gene is expressed in various tissues with tissue-specific alternative splicing (Honda, Harada, & Takagi, 1994; Mahendroo, Mendelson, & Simpson, 1993; Means, Kilgore, Mahendroo, Mendelson, & Simpson, 1991; Simpson et al., 1993). Patients harboring a loss-of-function mutation in the *CYP19A1* gene show severe osteoporosis and greater stature due to delayed or unsuccessful growth plate closure (Morishima, Grumbach, Simpson, Fisher, & Qin, 1995), both of which are sensitive to estrogen replacement therapy (Bilezikian, Morishima, Bell, & Grumbach, 1998; Carani et al., 1997; Rochira & Carani, 2009). Also, conventional aromatase knockout mice (ArKO) exhibited bone loss caused by decreased bone formation (Oz et al., 2001) or increased bone resorption (Miyaura et al., 2001). These reports indicate that the functions of aromatase are essential for the preservation of bone health. However, it is still uncertain whether a deficiency of aromatase alters bone metabolism directly or indirectly via systemic influences because both patients and ArKO mice show systemic endocrine disturbances.

Aromatase inhibitors (AIs) are applied clinically to treat certain populations of breast cancers in postmenopausal women (Goss et al., 2003). Although the serum levels of estrogen should be remarkably decreased in postmenopausal women, AIs still have adverse effects by increasing the risk of osteoporosis, called cancer treatment-induced bone loss (Becker et al., 2012). Furthermore, Finkelstein et al. reported that the administration of AIs to healthy men decreased bone mass in a serum testosterone concentration-independent manner (Finkelstein et al., 2016). These results suggest that part of the anabolic effect of testosterone (T) in men is dependent on E2 biosynthesis by aromatase. This evidence may indicate that the activities of aromatase in peripheral tissues and organs as well as the gonads are significant. Moreover, local synthesis of estrogens by aromatase may exert a variety of biological functions. However, it remains unclear how estrogens that are biosynthesized locally through the action of aromatase in non-gonadal tissues, contributes to the maintenance of bone mass.

In this study, we generated functional *Cyp19a1* floxed mice and osteoblast- or adipocyte-specific aromatase KO mice. Male adipocyte-specific aromatase KO mice (*Aro*^Δ*aP2*^) exhibited bone loss due to decreased serum phosphate and urinary phosphate reabsorption as a result of reduced NaPi2 expression in kidney. These results indicate that locally synthesized estrogens in adipocytes can play a significant role in bone mass maintenance. It also suggests the possible existence of a new phosphate regulatory mechanism through aromatase in adipose tissue.

## Results

### Generation of *Cyp19a1* floxed mice

To assess the tissue-specific function of aromatase, which is coded by the *Cyp19a1* gene, we generated *Cyp19a1* floxed mice. Figure S1 A shows the strategy used to generate the mutant *Cyp19a1* allele. Two loxPs were designed to be inserted at sites flanking exon 8 and exon 9 of the *Cyp19a1* allele as those exons correspond to the catalytic domain of aromatase (Ghosh, Griswold, Erman, & Pangborn, 2009). Successful insertion of NeoR cassette and loxPs was confirmed by Southern blotting using two different probes (Figure S1 B). Cre-mediated recombination excised exon 8, exon 9 and the NeoR gene from the mutant allele, resulting in the null allele. As expected, the *Cyp19a1* transcript was not detected in systemic *Cyp19a1* knockout (*Aro^-/-^*) mice by either qualitative or quantitative RT-PCR (Figure S1 C).

### Phenotypes of systemic and osteoblast-specific *Cyp19a1* knockout mice

To further confirm the abnormal phenotypes of the systemically expressed Cre-mediated deletion in *Cyp19a1* floxed mice, we determined whether *Aro^-/-^* mice showed the same phenotypic abnormalities reported in conventional *Cyp19a1* knockout mice (Fisher, Graves, Parlow, & Simpson, 1998; Jones et al., 2000). As expected, female *Aro^-/-^* mice exhibited smaller atrophic uteri and male *Aro^-/-^* mice showed significantly larger seminal vesicles (Figure S1 D, E). Also, *Aro^-/-^* mice showed a significantly increased volume of fat tissue, regardless of sex (Figure S1 E). These results suggested that aromatase was functionally depleted in *Cyp19a1* floxed mice when excised by Cre-mediated recombination of the *Cyp19a1* gene locus, as these phenotypes were similar to those in previous reports.

Next, we tried to reveal the function of aromatase in osteoblast lineage cells, because some previous studies reported that aromatase is expressed in osteoblasts (Sasano et al., 1997; Shozu & Simpson, 1998; Yanase et al., 2003). We generated three lines of osteoblast/osteocyte-specific *Cyp19a1* knockout mice by crossing *Cyp19a1* floxed (*Aro^flox/flox^*) mice with *2.3kb-Col1a1-Cre* (*Aro*^Δ*Col1a1*^), *OCN-Cre* (*Aro*^Δ*OCN*^), or *Dmp1-Cre* (*Aro*^Δ*Dmp1*^) mice. Bone mineral density (BMD) in *Aro*^Δ*Col1a1*^ female mice and *Aro*^Δ*OCN*^ male mice was comparable with controls (Figure S2 A, B). In contrast, BMD of *Aro*^Δ*Dmp1*^ mice was significantly greater in female but not male mice (Figure S2 C). However, we did not analyze this phenotype in detail because our goal was to determine why aromatase deficiency causes bone loss. Taken together, these results suggest that local aromatase expression in osteoblast/osteocyte lineage cells either does not contribute to bone mass maintenance or has a negative effect on bone mass.

### Deletion of *Cyp19a1* from adipose tissue resulted in decreased bone mass in male mice

Adipose tissue is known as the major site of estrogen biosynthesis or restoration in postmenopausal women and old men. Previous studies reported that aromatase expression in adipose tissue was increased by age, obesity, and menopause (Agarwal, Ashanullah, Simpson, & Bulun, 1997; Brown et al., 2017; Bulun & Simpson, 1994; Mair, Gaw, & MacLean, 2020). To investigate the effects of expressed aromatase in adipose tissue on bone, we generated adipocyte-specific *Cyp19a1* knockout mice (*Aro*^Δ*aP2*^) by crossing *Aro^flox/flox^* mice with *aP2-Cre* mice. *Cyp19a1* expression levels in epididymal white adipose tissue (eWAT) were significantly lower in *Aro*^Δ*aP2*^ compared to controls (Figure 1 A). In subcutaneous WAT (sWAT) and brown adipose tissue (BAT), *Cyp19a1* expression levels were quite low or undetectable (Figure S3 A). Although there were no differences in body weight (Figure 1 B), tibial and femoral BMD of *Aro*^Δ*aP2*^ male mice at the age of 16-20 weeks were significantly lower compared to control male but not female mice (Figure 1 C and Figure S2 D). Especially, proximal site BMD in the tibia was significantly less in *Aro*^Δ*aP2*^ mice compared to controls. The reason for the absence of the phenotype in females may be that *Aro*^Δ*aP2*^ female mice had sufficient estrogen production from the ovary and sufficient estrogen level in serum. Since T and E2 are involved in the maintenance of muscle mass and function (Sakakibara et al., 2021) (Sakai et al., 2024) (Kitajima & Ono, 2016), and administration of AIs has also been reported to alter body composition (Finkelstein et al., 2013), muscle and fat weight were measured. *Aro*^Δ*aP2*^ mice had low gastrocnemius muscle weight, and high eWAT weight compared to controls (Figure 1 D, E). These phenotypes of low bone and high fat mass were also observed in *Aro*^Δ*aP2*^ mice at the age of 24 weeks (Figure S4 A-E). It has been reported that some Cre mouse lines develop skeletal dysplasia or abnormal regeneration in tissues (Rodda & McMahon, 2006; Sakai et al., 2022). Therefore, we tested whether these phenotypes of *Aro*^Δ*aP2*^ mice were due to the influence of the Cre mouse line. The results showed that *aP2-Cre* mice were comparable in tibial and femoral BMD, fat mass, and muscle mass to littermate WT control mice (Figure S5 A-D). Also, to confirm the tissue-specificity of the *aP2-Cre* knockout approach, we checked *Cyp19a1* expression levels in bone, kidney, and muscle in *Aro*^Δ*aP2*^ and *Aro^flox/flox^* mice (Figure S3 A). No *Cyp19a1* expression was detected in femur, kidney and gastrocnemius muscle. To confirm aromatase activity, we also checked *Esr1* expression in these tissues; *Esr1* expression did not differ between *Aro*^Δ*aP2*^ and *Aro^flox/flox^* mice (Figure S3 B). We measured adipocyte size in eWAT and performed intraperitoneal glucose tolerance tests (IPGTT) because *Aro*^Δ*aP2*^ mice were normal-weight obesity (Franco, Morais, & Cominetti, 2016)-like phenotype. Adipocyte size was not different between groups (Figure 1 F). Fasting blood glucose was significantly higher overall in *Aro*^Δ*aP2*^ compared to controls, but there was no difference at any specific time point of IPGTT between groups (Figure 1 G). Since *Aro*^Δ*aP2*^ mice have suppressed E2 biosynthesis from T by aromatase in adipose tissue, serum T and E2 levels were measured by LC-MS/MS. There were no differences in serum E2 and T levels between groups (Figure 1 H). In addition, seminal vesicle weight in males and uterus weight in females were not different between *Aro*^Δ*aP2*^ and *Aro^flox/flox^* mice at 24 weeks old (Figure S6 A-C). These results suggest that E2 biosynthesized by aromatase in adipocytes works only locally, not systemically.

**Figure 1.**
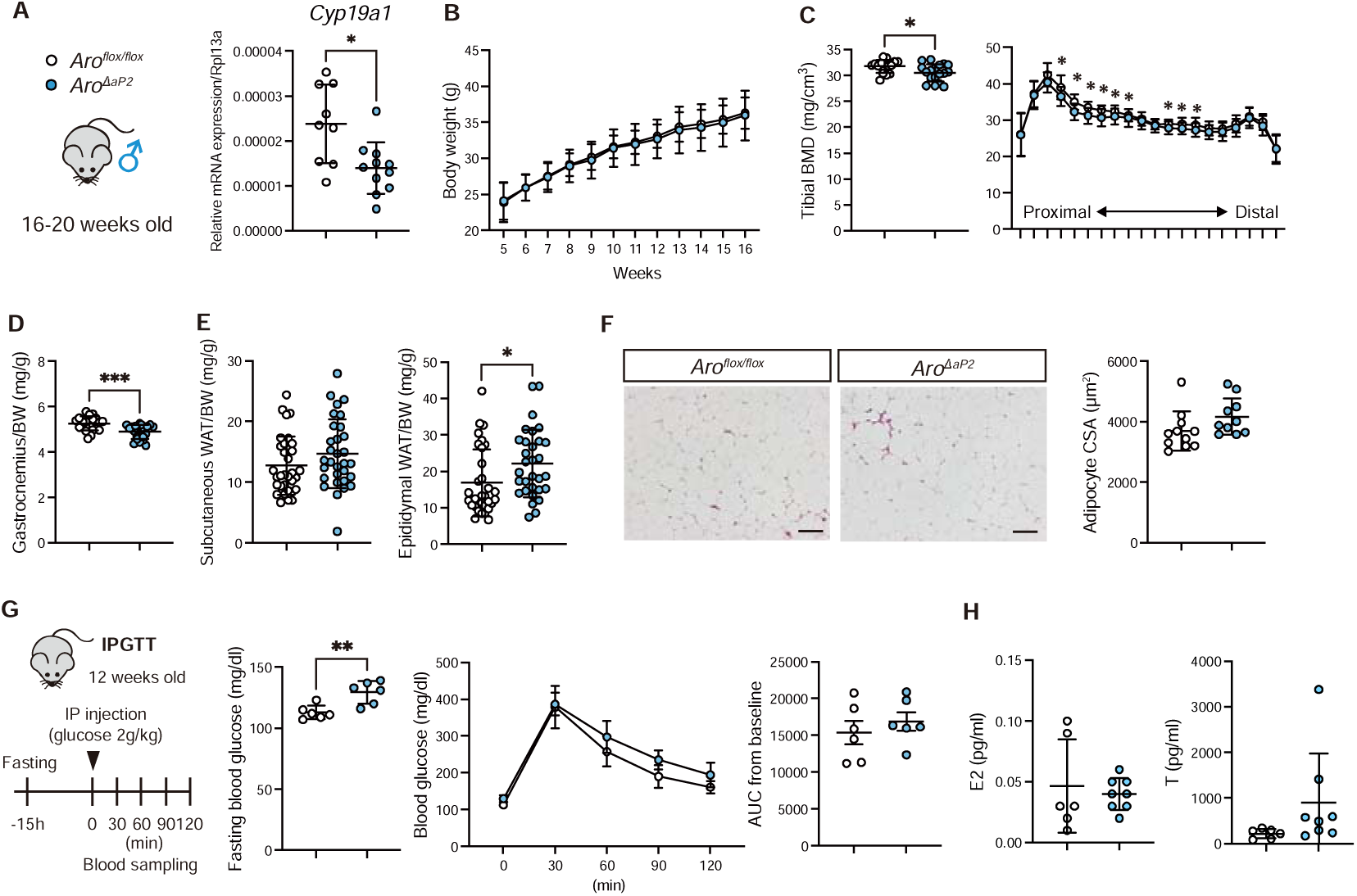
Male *Aro*^Δ*aP2*^ mice exhibited low bone mass, low muscle mass, high fat mass, and high fasting blood glucose. (A) *Cyp19a1* gene expression in eWAT. (B) Body weight. (C) Total tibial BMD (left) and BMD at each site when the bone is divided into 20 sections (right). (D) Gastrocnemius muscle weight per body weight. (E) Subcutaneous and epidydimal white adipose tissue weight per body weight. (F) Representative image and quantitative data of adipocyte cross sectional area (CSA) in eWAT. Scale bar indicates 100 µm. (G) Intraperitoneal glucose tolerance test (IPGTT) of 12 weeks old *Aro*^Δ*aP2*^ and *Aro^flox/flox^*mice. (H) Serum estradiol (E2) and testosterone (T) by LC-MS/MS. Data are presented as means ± SD. *, *p* < 0.05; **, *p* < 0.01; ***, *p* < 0.001

As E2 biosynthesis in adipocytes is reduced due to aromatase ablation in *Aro*^Δ*aP2*^, it is presumed that E2/estrogen receptor (ER) signaling is also reduced. In fact, we examined steroid hormone receptor expression in eWAT and found that *Esr1* expression was significantly less in *Aro*^Δ*aP2*^ compared to control mice (Figure S7 A). Therefore, to investigate whether the *Aro*^Δ*aP2*^ phenotype is due to local ERα action in adipocytes, we generated adipose tissue-specific *Esr1* knockout mice (*ER*α^Δ*aP2*^) by crossing *ER ^flox/flox^* mice with *aP2-Cre* mice. *Esr1* expression levels in eWAT were significantly lower in α compared to control mice (Figure S7 B). The results showed no group differences in body weight, bone or muscle mass (Figure S7 C, D, F). In contrast, subcutaneous WAT (sWAT) and eWAT weight were significantly greater in *ER*α^Δ*aP2*^ compared to control mice (Figure S7 E). This phenotype was similar to that of *Aro*^Δ*aP2*^ mice. These results suggest that ERα in adipocytes is involved in lipid metabolism but not bone metabolism. Therefore, the bone phenotype in *Aro*^Δ*aP2*^ was not an ERα-mediated effect of adipocytes.

### *Aro*^Δ^*^aP2^* mice had an abnormal phosphate metabolism

We analyzed bone structure and bone marrow fat volume of the proximal tibia of *Aro*^Δ*aP2*^ and *Aro^flox/flox^*mice using µCT. Trabecular bone parameters such as trabecular bone volume (BV/TV) and trabecular thickness (Tb.Th) were significantly lower in *Aro*^Δ*aP2*^ compared to control mice (Figure 2 A). However, cortical bone parameters, and regulated bone marrow tissue (rBMAT) volume were not different between groups (Figure 2 B, C). Further, serum P1NP, a bone formation marker, and serum CTX, a bone resorption marker, were not different between groups (Figure 2 D).

**Figure 2.**
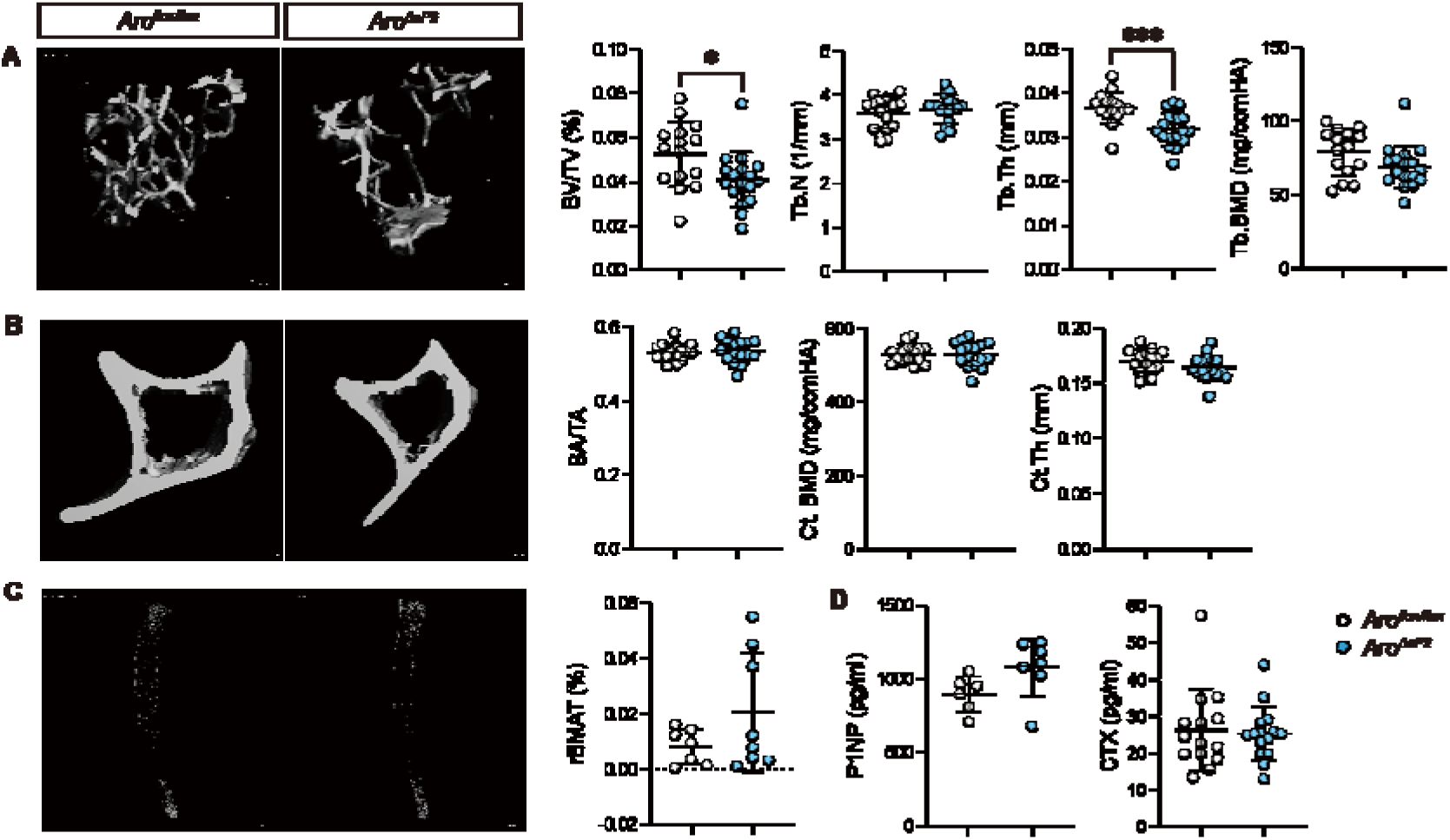
*Aro*^Δ*aP2*^ mice exhibited low trabecular bone volume at the proximal tibia. Representative image and quantitative data of (A) bone volume per tissue volume (BV/TV), trabecular number (Tb.N), trabecular thickness (Tb.Th), and trabecular bone mineral density (Tb.BMD) as trabecular bone parameters, (B) bone area per tissue area (BA/TA), cortical bone mineral density (Ct.BMD), and cortical thickness (Ct.Th) as cortical bone parameters, and (C) regulated bone marrow tissue (rBMAT) volume at the proximal tibia. (D) Serum P1NP and CTX concentration. Data are presented as means ± SD. *, *p* < 0.05; ***, *p* < 0.001

To investigate the dynamics of osteoclasts and osteoblasts, we performed bone histomorphometry in the proximal tibia of *Aro*^Δ*aP2*^ and *Aro^flox/flox^* mice. von Kossa/van Gieson staining revealed significantly greater osteoid volume (OV/BV) and osteoid thickness (O.Th) in *Aro*^Δ*aP2*^ compared to control mice (Figure 3 A, B). In addition, TRAP staining revealed significantly lower osteoclast surface/bone surface (Oc.S/BS) and osteoclast number/bone perimeter (N.Oc/B.Pm) in *Aro*^Δ*aP2*^ compared to control mice (Figure 3 C, D). However, bone formation rate and osteoblast number were not different between groups (Figure 3 E-H). These results suggest that *Aro*^Δ*aP2*^ mice exhibit an abnormal bone calcification phenotype.

**Figure 3.**
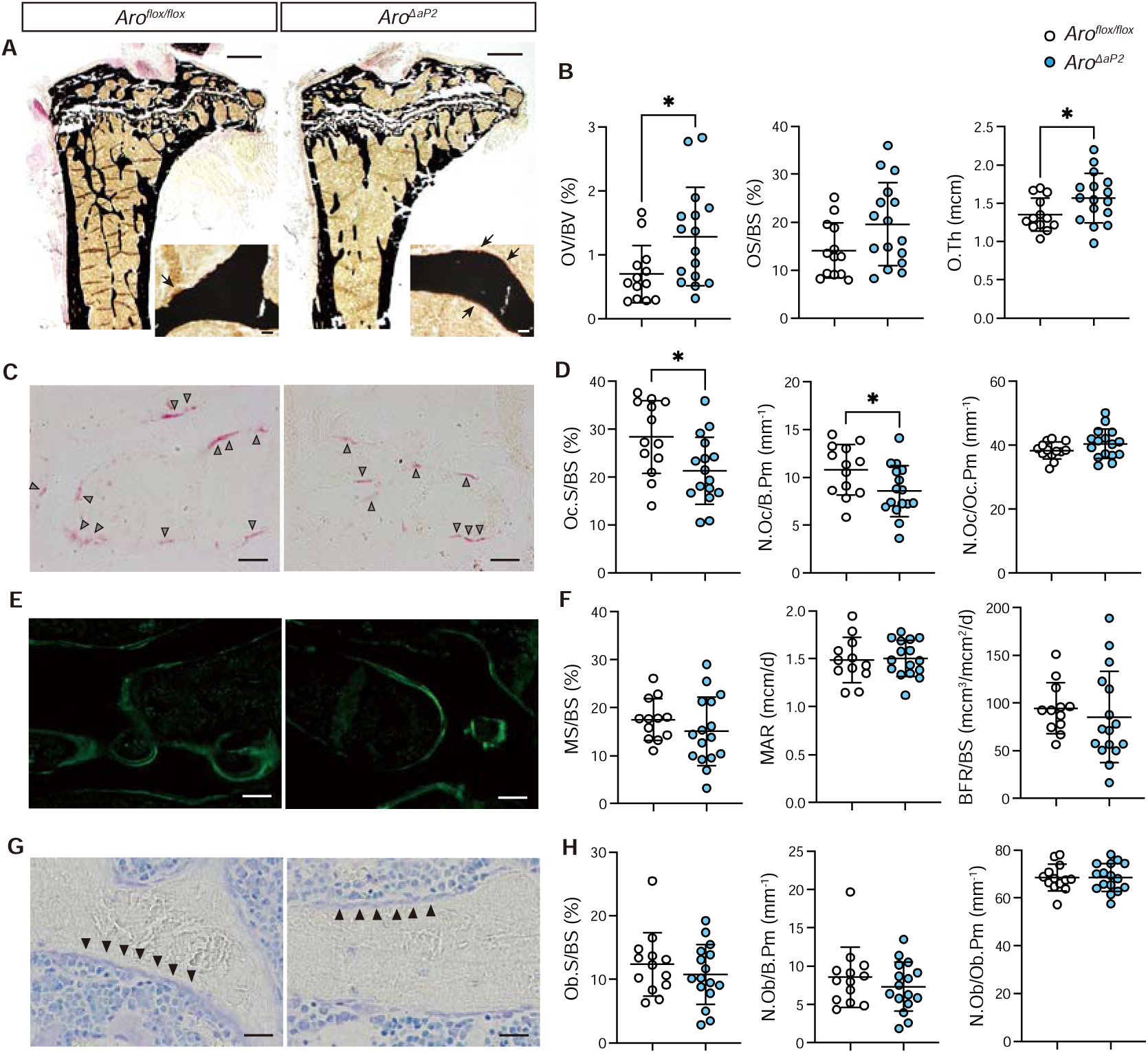
*Aro*^Δ*aP2*^ mice exhibited high osteoid volume and low osteoclast number. (A) Representative image of von Kossa/van Gieson staining in the tibia. Black arrow indicates osteoid. Scale bar of low magnification image indicates 500 µm. Scale bar of high magnification image indicates 20 µm. (B) Osteoid volume/bone volume (OV/BV), osteoid surface/bone surface (OS/BS), osteoid thickness (O.Th). (C) Representative image of TRAP staining in tibia. Gray arrowhead indicates osteoclast. Scale bar indicates 50 µm. (D) Osteoclast surface/bone surface (Oc.S/BS), osteoclast number/bone perimeter (N.Oc/B.Pm), osteoclast number/osteoclast perimeter (N.Oc/Oc.Pm). (E) Representative image of calcein labeling in tibia. Scale bar indicates 100 µm. (F) Mineralizing surface (MS/BS), Mineral apposition rate (MAR), Bone formation rate (BFR/BS). (G) Representative image of Toluidine blue staining in tibia. Black arrowhead indicates osteoblast. Scale bar indicate 50 µm. (H) Osteoblast surface (Ob.S/BS), osteoblast number/bone perimeter (N.Ob/B.Pm), osteoblast number/osteoblast perimeter (N.Ob/Ob.Pm). Data are presented as means ± SD. *, *p* < 0.05

Dysregulated phosphate metabolism is an important cause for insufficient calcification. To assess this, we analyzed serum and urine parameters related to phosphate metabolism of *Aro*^Δ*aP2*^ and *Aro^flox/flox^* mice. Serum inorganic phosphate (IP) was significantly lower in *Aro*^Δ*aP2*^ mice compared to control mice but not in urine (Figure 4 A, B). Other parameters in serum and urine such as calcium (Ca), alkaline phosphatase (ALP), and creatinine (CRE) were not different between groups (Figure 4 A, B). Then we calculated urinary phosphate reabsorption using serum and urine parameters. Tmp/GFR, an indicator of phosphate reabsorption in the renal proximal tubules was significantly lower in *Aro*^Δ*aP2*^ compared to control mice (Figure 4 C). In addition, serum IP and OV/BV were significantly and negatively correlated (Figure 4 D). These results suggest that *Aro*^Δ*aP2*^ mice have high osteoid volume and low bone mass due to decreased serum IP caused by decreased urinary phosphate reabsorption.

**Figure 4.**
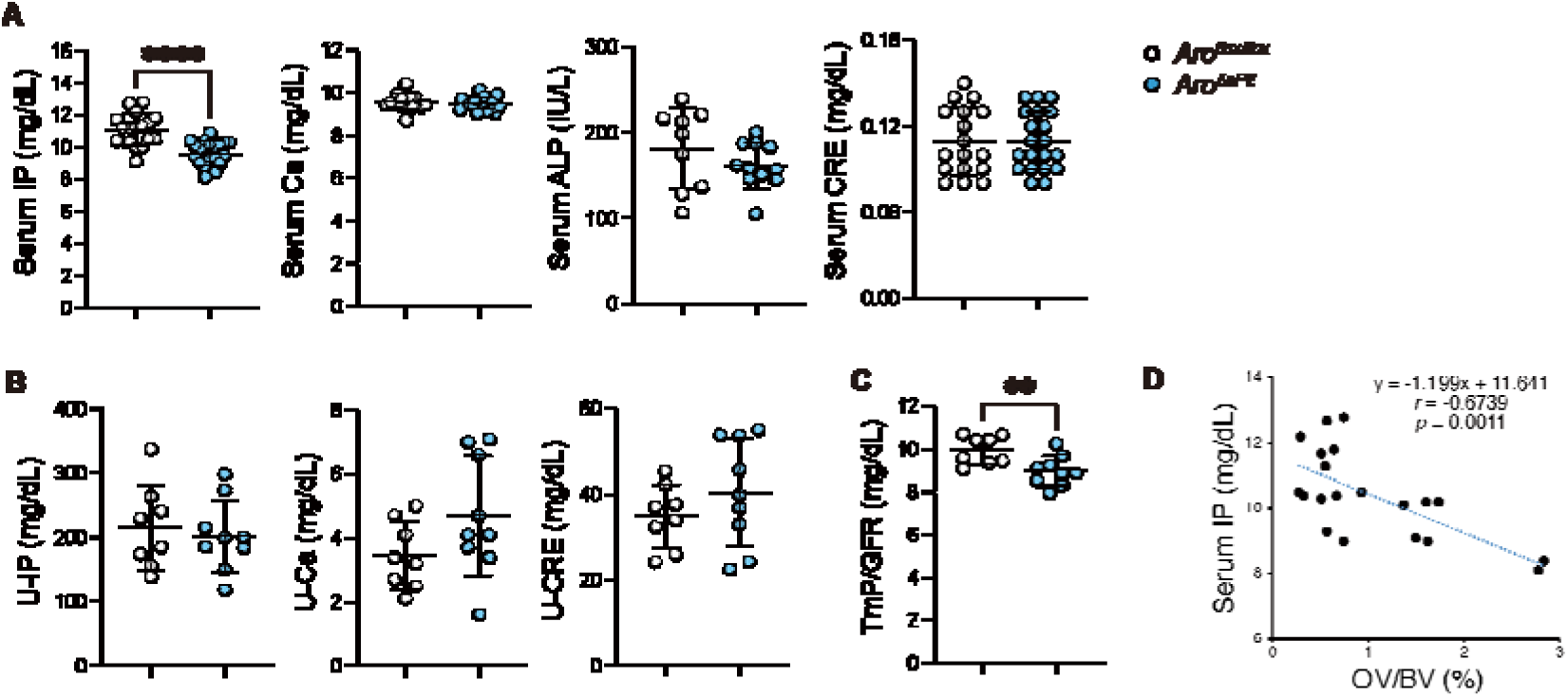
*Aro*^Δ*aP2*^ mice exhibited low serum phosphate and decreased urinary phosphate reabsorption. (A) Serum IP, Ca, ALP and CRE. (B) Urine IP, Ca, and CRE. (C) Tmp/GFR calculated by serum and urine parameters. (D) Correlation between serum IP and OV/BV. Data are presented as means ± SD. **, *p* < 0.01; ****, *p* < 0.0001.

To identify the responsible genes for the phenotypes due to lack of *Cyp19a1* in adipose tissue that regulate phosphate metabolism, we performed RNA-seq analysis using total RNA from eWAT in *Aro*^Δ*aP2*^ and *Aro^flox/flox^* mice. The PCA plot showed separation between the *Aro*^Δ*aP2*^ and *Aro^flox/flox^*(Figure S8 A). The results yielded 125 genes with a significant increase and 100 genes with a significant decrease (*p* < 0.01) (Figure S8 B). We focused on the genes that showed significant changes and performed Gene Ontology (GO) analysis. Genes involved in inflammatory responses were increased and genes involved in nucleotide metabolic processes were decreased in *Aro*^Δ*aP2*^ compared to controls (Figure S8 C, D), suggesting that the adipose tissue of *Aro*^Δ*aP2*^ mice was in an inflammatory state and metabolically compromised. However, there were no genes encoding proteins that act on other organs as a humoral factor among differentially expressed genes.

Therefore, we decided to focus our analyses on existing and known factors that can regulate phosphate metabolism. Phosphate reabsorption occurs in the proximal tubule cells of the kidney, mainly via Na^+^-Pi cotransporter 2a (NaPi2a) and Na^+^-Pi cotransporter 2c (NaPi2c). Phosphate homeostasis is under endocrine control that is mediated by FGF23, PTH and 1,25LJdihydroxyvitamin D (1α,25(OH)_2_D_3_) (Fukumoto & Martin, 2009; Minisola et al., 2017). Of these, FGF23, PTH, and corticosterone suppress NaPi2 expression or function in the renal tubule (Walton & Bijvoet, 1975). To investigate why *Aro*^Δ*aP2*^ mice have low serum phosphate levels, we analyzed several parameters that can be influenced by proximal tubular function and NaPi2 expression levels in the kidney. Urine beta-2-microglobulin (B2M) and serum uric acid (UA), indicators for renal tubular damage, and Na, K, and CL, indicators for Fanconi syndrome, were not different between groups (Figure 5 A-C). In addition, serum PTH, 1α,25(OH)_2_D_3_, 25(OH)D_3_, and corticosterone were not different between groups (Figure 5 D, E, G). In contrast, serum FGF23 levels were significantly lower in *Aro*^Δ*aP2*^ compared to control mice (Figure 5 F). Initially we hypothesized that high FGF23, as seen in patients with FGF23-related hypophosphatemic rickets/osteomalacia, was the cause of low phosphate in *Aro*^Δ*aP2*^ mice. However, serum FGF23 probably decreased in response to lower serum phosphate concentrations, suggesting that the insufficient calcification phenotype in *Aro*^Δ*aP2*^ mice was not due to increased FGF23. Next, we analyzed NaPi2 expression in the kidney. Transcript levels of *Npt2a* and *Npt2c* in the kidney were not different between groups (Figure 5 H). As NaPi2 activity can be regulated by endocytosis and membrane recycling of NaPi2 protein in addition to transcriptional and translational regulations, protein levels of NaPi2a and NaPi2c in brush border membrane vesicles (BBMVs) were confirmed by western blotting. Protein levels of NaPi2a and NaPi2c in BBMVs were significantly decreased in *Aro*^Δ*aP2*^ compared to control mice (Figure 5 I). In *Aro*^Δ*aP2*^ mice, phosphate-regulating hormones such as PTH, 1α,25(OH)_2_D_3_, and corticosterone, previously reported to regulate NaPi2 in the kidney, were normal and serum FGF23 was lower. Taken together, these results indicate that renal NaPi2 is decreased in *Aro*^Δ*aP2*^ mice, independent of known NaPi2 regulation mechanisms.

**Figure 5.**
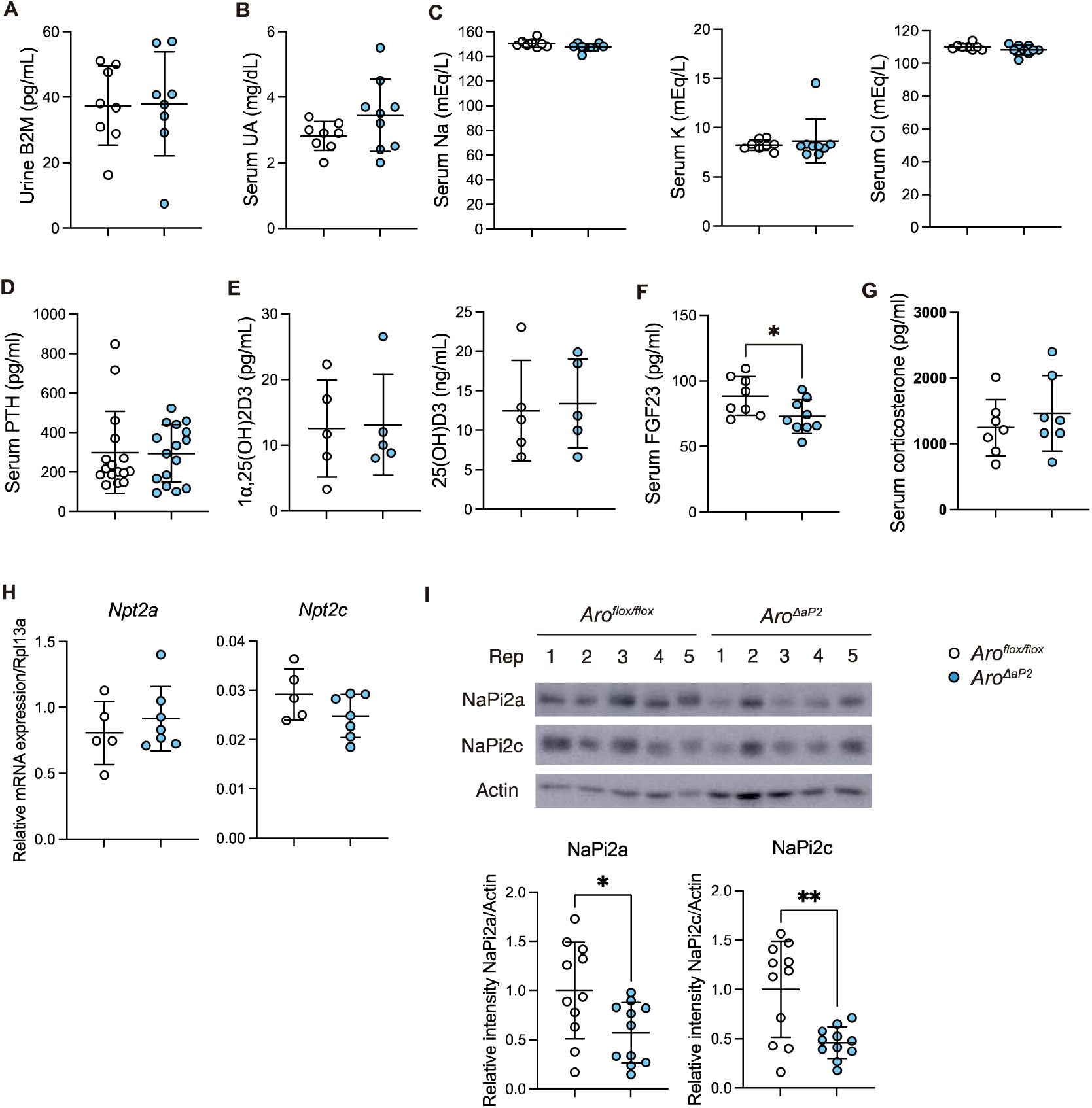
*Aro*^Δ*aP2*^ mice exhibited low NaPi2a and NaPi2c protein levels in the kidney. (A) Urine B2M and (B) Serum UA as indicators for renal tubular damage. (C) Serum Na, K, and Cl as indicators for Fanconi syndrome. (D) Serum PTH, (E) 1α,25(OH)_2_D_3_, 25(OH)D_3_, (F) FGF23, and (G) corticosterone as a phosphate metabolism regulating hormone. (H) Gene expression of *Npt2a* and *Npt2c* in the kidney. (I) Protein expression of NaPi2a and NaPi2c in renal BBMVs. Western blotting images showed 5 samples out of 11samples for each group. Data are presented as means ± SD. *, *p* < 0.05; **, *p* < 0.01.

To determine whether low BMD in *Aro*^Δ*aP2*^ mice could be ameliorated by additional phosphate intake, we performed an experiment in which *Aro*^Δ*aP2*^ mice were fed with high-phosphate water for 4 weeks (Figure S9 A). However, tibial BMD was not different between high phosphate water (HP) and tap water (CON) condition (Figure S9 C). Serum IP, Ca, ALP, Urine IP, CRE, Ca, and TmP/GFR were also not different between both conditions (Figure S9 D-F). On the other hand, serum CRE levels were significantly higher in HP condition compared to CON (Figure S9 D), suggesting HP may worsen renal function in *Aro*^Δ*aP2*^ mice. Therefore, since *Aro*^Δ*aP2*^ mice have reduced phosphorus reabsorption in the renal tubules due to decreased expression of NaPi2a and NaPi2c, HP could not rescue BMD in *Aro*^Δ*aP2*^ mice and may further worsen renal function.

## Discussion

In this study, we investigated the osteoprotective function of aromatase in the non-gonadal tissues, bone and adipose. Although three different types of osteoblast/osteocyte-specific aromatase KO mice exhibited no low bone mass phenotype, *Aro*^Δ*aP2*^ male mice but not female mice exhibited abnormal bone calcification with low serum phosphate levels due to lower urinary phosphate reabsorption caused by decreased NaPi2a and NaPi2c in the kidney (Figure 6).

**Figure 6.**
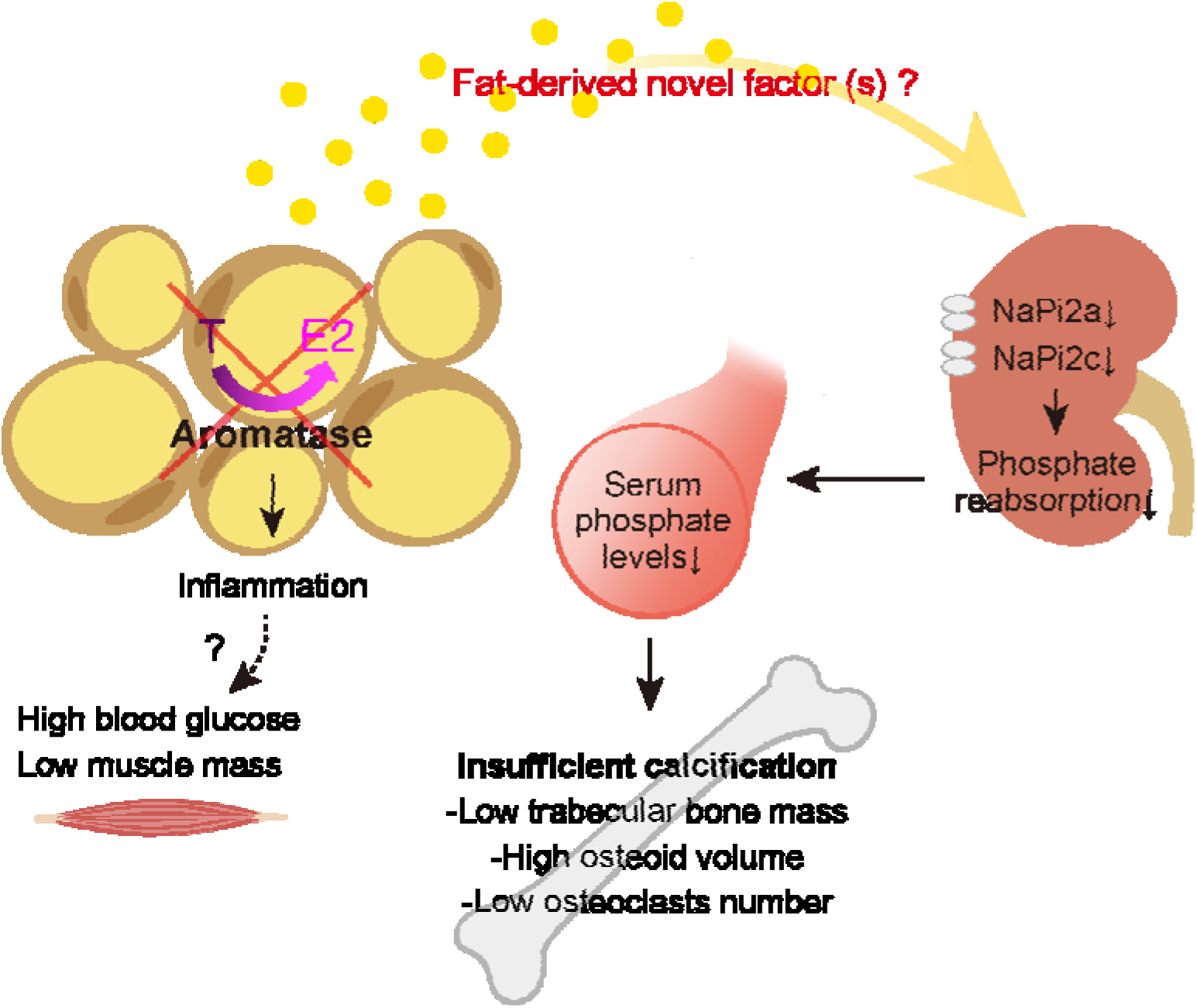
In *Aro*^Δ*aP2*^ mice, E2 biosynthesis by aromatase in adipose tissue was suppressed, resulting in suppression of fat-derived novel factor (s) that regulates renal NaPi2 protein levels, suppressing renal phosphate reabsorption and reducing serum phosphate levels. As a result, *Aro*^Δ*aP2*^ mice exhibited an abnormal bone calcification phenotype.

The *Aro*^Δ*aP2*^ mice show an increase in osteoid and a decrease in osteoclast, which is similar to the phenotype of osteomalacia (Eicher, Southard, Scriver, & Glorieux, 1976; Hayashibara et al., 2007), suggesting that an insufficient calcification was occurring. However, the *Aro*^Δ*aP2*^ mice do not exhibit abnormal bone phenotype as large as osteomalacia, and their serum IP levels are only slightly decreased, so we believe that they have an insufficient calcification that does not affect osteoblast numbers. In addition, since *ER* ^Δ*aP2*^ mice not showed low bone mass, non-genomic or classical effects of ERα in adipocytes may not contribute to bone phenotype in *Aro*^Δ*aP2*^ mice. White adipose tissue contained many cells such as adipocytes, preadipocytes, endothelial cells, and immune cells (Emont et al., 2022). Thus, we speculate that synthesized estrogen by aromatase in adipocytes affect not only adipocytes itself but also other cell types in adipose tissue, then maintaining bone mass via unknown factors. *Aro*^Δ*aP2*^ mice exhibited not only bone loss but also low muscle mass, high fat mass, and high fasting blood glucose, partially similar to symptoms of late-onset hypogonadism (LOH), an age-related androgen deficiency (Lunenfeld, Saad, & Hoesl, 2005). Furthermore, Finkelstein et al. examined changes in body composition (Finkelstein et al., 2013) and bone mass (Finkelstein et al., 2016) in healthy men aged 20-50 years when the aromatization of T to E2 was inhibited using goserelin acetate which suppresses endogenous gonadal steroid production, testosterone gel, and AIs such as anastrozole. The results showed that inhibition of aromatization of T to E2 by anastrozole increased the percentage of body fat (Finkelstein et al., 2013) and decreased trabecular BMD as measured by quantitative computed tomography (Finkelstein et al., 2016). These findings suggest that the effects of T are dependent on a predominantly estrogenic effect for fat and bone in men. Furthermore, it has been reported that mice overexpressing aromatase in white adipose tissue using aP2 promoter-hCYP19 fusion gene (*Ap2-arom* mice) have shown improved insulin sensitivity and reduced fat inflammation in male mice. Further, the *Ap2-arom* mice exhibited decreasing the expression of markers of macrophages and immune cell infiltration in WAT of male (Ohlsson et al., 2017). This previous study supported the biological significance of aromatase in adipose tissue as same as our study because the phenotypes including RNA-seq and blood glucose data of *Ap2-arom* mice was completely opposite to our *Aro*^Δ*aP2*^ mice, while there was no report about bone phenotype of *Ap2-arom* mice. As the phenotype of *Aro*^Δ*aP2*^ mice was similar to the results of these studies, estrogen, which is synthesized by aromatase in fat, could play an important role in maintaining body composition as well as bone in men.

On the other hand, *Aro*^Δ*aP2*^ mice also exhibited low muscle mass, which may be caused by fat inflammation. Previously reported that obesity activate macrophages, mast cells and T lymphocytes, promoting a low-level inflammation that results in the secretion of pro-inflammatory cytokines such as tumour necrosis factor (TNF) and IL-6, then these secretory changes lead to muscle catabolism and insulin resistance, promoting gain in fat mass and a loss of muscle mass (Batsis & Villareal, 2018). According to our GO analysis in RNA-seq, gene expressions of adipose tissue in *Aro*^Δ*aP2*^ mice were enriched fat inflammation related term and macrophage related genes. Therefore, it is possible that fat inflammation in *Aro*^Δ*aP2*^ mice induced muscle catabolism and insulin resistance, leading to muscle loss and elevated fasting blood glucose. However, we did not perform study in detail in this point of view, thus we need further study.

The present study suggested that *Aro*^Δ*aP2*^ mice have lower serum phosphate levels due to lower urinary phosphate reabsorption caused by decreased NaPi2a and NaPi2c in the kidney. Phosphate homeostasis is strictly regulated by hormones such as FGF23, PTH and 1,25(OH)_2_D_3_. Among them, FGF23 was established as the principal hormone in regulating blood phosphate level (Shimada et al., 2004). As a mechanism, FGF receptor type 1 (FGFR1) works as a phosphate-sensing receptor, and its downstream intracellular signaling pathway regulates the expression of *GALNT3*. And then, the post-translational modifications of FGF23 protein via a gene product of *GALNT3* enhance FGF23 production. As results, FGF23 can reduce blood phosphate level by inhibiting both proximal tubular phosphate reabsorption and intestinal phosphate absorption via downregulating the activity of NaPi2a and NaPi2c in the kidney and blood 1α,25(OH)_2_D_3_ level (Takashi, 2024; Takashi et al., 2019; Takashi et al., 2021). However, *Aro*^Δ*aP2*^ mice has normal phosphate-regulating hormones’ level and low serum FGF23. Therefore, the existence of novel control mechanisms other than those mentioned factors above was suggested.

There are some limitations in this study. First, we tried to measure E2 level in fat tissue by LC-MS/MS, but it could not be detected. Since ER signaling in eWAT was significantly lower in *Aro*^Δ*aP2*^ compared to control mice, E2 was expected to be lower in *Aro*^Δ*aP2*^. Second, we did not analyze patient data. There are no reports of low serum phosphate levels in AI-treated patients or patients with loss-of-function mutations of the *Cyp19a1* gene. However, as the phenotype in the present study was mild, it is expected that the decrease in phosphate in patients might be a mild decrease within the reference range. This requires further investigation. Third, we did not analyze *Aro*^Δ*aP2*^ female mice in detail. It is hypothesized that the intact *Aro*^Δ*aP2*^ female mice had sufficient estrogen production from the ovaries and thus had no phenotype (Figure S2 D). Therefore, it will be necessary to analyze *Aro*^Δ*aP2*^ mice in which estrogen deficiency is induced by ovariectomy or other means. Fourth, although we generated and analyzed bone- and adipose tissue-specific aromatase knockout mice, the brain is also known to highly express aromatase and may contribute to skeletal homeostasis (Kim et al., 2021). Therefore, future studies are needed to examine the function of aromatase in other tissues, including brain, to bone homeostasis. Finally, in this study, we were unable to identify a novel factor that regulates NaPi2 protein levels. All known predicted regulators were examined, suggesting the existence of a novel phosphoregulatory mechanism that has not been previously identified.

In conclusion, we have shown that E2 biosynthesis by aromatase in fat plays an important role in the maintenance of bone mass in male via phosphate metabolism.

## Materials and Methods

### Generation of *Cyp19a1* floxed mice by gene targeting

The PL451 vector, which contains a neo cassette flanked by two frt sites and one loxP site, was obtained from NCI-Frederick. A cassette vector was generated by inserting the second loxP site and the DT-A gene from the pMC1DTpA vector into PL451. Three fragments containing the mouse *Cyp19a1* gene were amplified by PCR and inserted into the cassette vector to form the final *Cyp19a1* knockout construct. The *Cyp19a1* conditional knockout targeting vector was linearized by PmeI and was introduced into M1 mouse embryonic cells (RIKEN) by electroporation and screened by genomic Southern blotting. Probes for Southern blotting was amplified by PCR using the following primers: Probe A, 5’-GGCCAGAGTCTGTCTGGAGT-3’, and 5’-AAGCTCTGGCAGGTGTCAAT-3’; Probe B, 5’-ACCCAAAAGCCTCAACACAG-3’, and 5’-GGCTTTCAGGACAAACCTTG-3’. Chimeric mice were generated by aggregation of ES with eight cell embryos of BDF1 mice. Chimeric mice were crossed with wild-type C57BL/6J mice (CLEA Japan) and the resulting pups with *Cyp19a1^flox/+^* were genotyped with the following primers: 5’-TAGGTGGCGAGAGAAAGGAA-3’, and 5’-CCCCAGAGGCTTACACACAT-3’ (WT allele; 411 bps, floxed allele; 482 bps).

### Generation of systemic and osteoblast- or adipocyte- specific knockout mice

*Cyp19a1^-/+^* mice were generated by crossing *Cyp19a1^flox/+^* mice with *CMV-Cre* mice (Dupé et al., 1997), which were kindly provided by Prof. Pierre Chambon. The null allele was genotyped with the following primers: 5’-ATTCCCGCCGAATAATCTCT-3’, and 5’-ACCTTCCTTGACCAATGTCG-3’ (Floxed [L2] allele; 6118 bps, KO [L-] allele; 315 bps). *2.3kb-Col1a1-Cre* transgenic mice, which were kindly provided by Dr. Gerard Karsenty (Dacquin, Starbuck, Schinke, & Karsenty, 2002), were crossed with *Cyp19a1^flox/flox^*to generate *Aro*^Δ*Col1a1*^ mice. *Osteocalcin-Cre* mice (Zhang et al., 2002), which were obtained from the Jackson Laboratory, were crossed with *Cyp19a1^floxflox+^* to generate *Aro*^Δ*OCN*^ mice. *Dmp1-Cre* transgenic mice, kindly provided by Prof. Lynda Bonewald (Lu et al., 2007), were crossed with *Cyp19a1^floxflox+^* to generate *Aro*^Δ*Dmp1*^ mice. *aP2-Cre* mice (He et al., 2003), which were obtained from the Jackson Laboratory (JAX stock #005069), were crossed with *Cyp19a1^flox/flox^* or *Esr1^flox/flox^*to generate *Aro*^Δ*aP2*^ or α mice. *Esr1* mice were kindly provided by Prof. Chambon (Dupont et al., 2000). Mice were housed in a specific pathogen-free facility under climate-controlled conditions at room temperature (22 ± 2°C) with 25% humidity and a 12-hr light/dark cycle. They were provided with ad libitum access to food and water. Animal experiments were approved by the Animal Experiment Committee of Ehime University (approval no. 37A1-1/16) and were performed in accordance with the Ehime University Guidelines for Animal Experiments.

### RT-qPCR

RNA was extracted from eWAT and kidney using a RNeasy Mini Kit (70146, QIAGEN) with QIAzol Lysis Reagent (79306, QIAGEN) or ISOGEN (319-90211, NIPPON GENE) and synthesized into cDNA using PrimeScript RT Master Mix (RR036, Takara Bio Inc.). RT-qPCR was performed using cDNA to evaluate gene expression with Thermal Cycler Dice (Takara Bio, Inc.) and SYBR Premix Ex Taq II (RR820S, Takara Bio Inc.). *Rpl13a* or *Gapdh* was used as the housekeeping gene. Primers are shown in Table S1.

### Radiological examination

Samples were fixed in 4% paraformaldehyde phosphate buffer solution (PFA) overnight at 4°C. The areal bone mineral density (aBMD) of the isolated tibiae and femora was measured by Dual-energy X-ray absorptiometry (DXA) using a bone mineral analyzer (DCS-600EX, ALOKA). Micro-computed tomography (μCT) scanning of the femora was performed according to the manufacturer’s instructions using a Scanco Medical μCT35 System (SCANCO Medical) with an isotropic voxel size of 6 μm. We defined the regions of interest (ROI) as 200 slices starting 1500 μm proximal to the proximal growth plate in the tibia. These images were used for 3D reconstruction and analysis. Structural parameters (3D) included cortical structure [bone area per total area (BA/TA), BMD, and cortical thickness (Ct.Th)], and trabecular structure [bone volume per total volume (BV/TV), trabecular number (Tb.N), trabecular thickness (Tb.Th), and BMD] according to established guidelines (Bouxsein et al., 2010).

### Osmium staining

Tibiae were harvested, fixed in 4% PFA overnight, and then immersed in 0.1 M EDTA (pH 7.3) for 2 weeks for decalcification. The decalcified tibiae were immersed in a 1:1 mixture of 5% potassium dichromate and 2% osmium tetroxide (23310-10, Polysciences, Inc., USA) for 48 h and stained. The stained tibiae were scanned using µCT to evaluate marrow fat volume (Scheller et al., 2014).

### Histology and histomorphometry

For bone histomorphometry, the mice were double labeled with subcutaneous injections of 15 mg/kg of calcein (Sigma) at 5 and 2 days before sacrifice. Tibiae were removed from each mouse and fixed with 4% PFA. After µCT analysis, tibiae were embedded with Methylmethacrylate (MMA) after dehydration and the plastic sections were cut by a standard microtome (LEICA) into 6 μm for von Kossa/van Gieson, TRAP, and Toluidine blue staining. The region of interest (ROI) was 750×1200 mm of secondary spongiosa in the tibia. Each section was photographed using a cellSense Ver. 1.3 (OLYMPUS) and analyzed using OsteoMeasure V3.3.0.1 (OsteoMetrics, Inc., GA, USA).

### Serum and urinary analysis

Blood samples were collected from heart when mice were sacrificed. Blood samples were centrifuged at 3,000 g at 4°C for 10 min, and separated serum samples were frozen and stored at -80°C until analysis. Urine samples were collected for 24 hours on the day before mice were sacrificed and stored at -80°C until analysis. Serum estradiol (E2), testosterone (T), 1,25(OH)_2_D_3_, and 25(OH)D_3_ were analyzed by LC-MS/MS. These analyses were outsourced to ASKA Pharma Medical Co., Ltd. Serum P1NP (MBS2702748, MyBioSource, Inc.), CTX (AC-06F1, Immunodiagnostic Systems), PTH (60-2305, QuidelOrtho Corporation.), intact FGF23 (TCY4000 (11A735), KAINOS Laboratories, Inc.), corticosterone (K014-H1/H5, Arbor Assays Inc.), and urine beta-2-microglobulin (ab223590, abcam) were analyzed by ELISA respectively. Analysis of serum IP, Ca, ALP, CRE, UA, Na, K, Cl, urine IP, Ca, and CRE were outsourced to Oriental Yeast Co., ltd.

### Protein sample purification and immunoblotting

BBMVs were prepared using the Ca^2+^ precipitation method. Cortical membrane and whole homogenate were obtained from mouse kidneys and used for immunoblotting analyses as described previously (Hanazaki et al., 2020; Ikuta et al., 2019; Ikuta et al., 2018; Kaneko et al., 2018). Protein samples were heated at 95°C for 5 min in sample buffer in the presence of 2-mercaptoethanol and subjected to sodium dodecyl sulfate-polyacrylamide gel electrophoresis (SDS-PAGE). The separated proteins were transferred by electrophoresis to Immobilon-P polyvinylidene difluoride membrane (Millipore, Billerica, MA, USA) and treated with diluted antibodies. Signals were detected using Immobilon Western (Millipore) (Hanazaki et al., 2020; Ikuta et al., 2019; Ikuta et al., 2018; Kaneko et al., 2018). Rabbit anti-NaPi2a and NaPi2c polyclonal antibodies were generated as described previously (Hanazaki et al., 2020; Segawa et al., 2009; Segawa et al., 2005; Segawa et al., 2007). Mouse anti-actin monoclonal antibody (MAB1501, Millipore, RRID: AB_2223041) was used as an internal control. Horseradish peroxidase-conjugated anti-rabbit IgG was utilized as the secondary antibody (115-035-003, Jackson ImmunoResearch Laboratories, Inc, West Grove, PA, USA, RRID: AB_10015289). The diluted antibodies for immunoblotting were: anti-NaPi2a (1:15,000), anti-NaPi2c (1:1500), and anti-actin (1:10,000).

### RNA-seq analysis

Total RNA was extracted from eWAT using RNeasy Mini Kit (70146, QIAGEN) with QIAzol Lysis Reagent (79306, QIAGEN). BRB-seq (Alpern et al., 2019) was performed for preparing libraries with some following modifications. Oligo-dT based primer was used for single-stranded synthesis and Second Strand Synthesis Module (NEB, #E6111) was used for double-stranded cDNA synthesis. In-house MEDS-B Tn5 transposase (Picelli et al., 2014; Sato et al., 2019) was used for tagmentation and amplified by 10 cycles of PCR using Phusion High-Fidelity DNA Polymerase (Thermo Scientific, #M0530). 51bp of insert read (Read2) were obtained on Illumina NovaSeq6000. Obtained FASTQ files were analyzed by RaNA-seq (Prieto & Barrios, 2020). Data were registered in GEO with the accession number GSE263849. Gene Ontology Analysis (GO) was conducted by DAVID (Huang da, Sherman, & Lempicki, 2009; Sherman et al., 2022) for significantly (*p* < 0.01) changed genes.

### Statistical analysis

The data are presented as means ± SD using unpaired t tests with Welch’s correction. The Brown-Forsythe and Welch ANOVA test was used for comparison among the four groups, and Dunnett T3 was used for post-hoc analysis. Outliers were excluded by SmirnovLJGrubbs test. All statistical analyses were performed using Graph Pad Prism Version 10.0.2 (GraphPad Software Inc., La Jolla, CA) and Microsoft Excel. P values less than 0.05 were considered significant.

## Acknowledgments

We are grateful to Dr. Erina Inoue for her technical support. We thank the staff of the Division of Analytical Bio-Medicine and the Division of Laboratory Animal Research, the Advanced Research Support Center (ADRES), the members of the Division of Integrative Pathophysiology, Proteo-Science Center (PROS), and Ehime University for their technical assistance and helpful support. We also thank the Medical Research Center for High Depth Omics, Medical Institute of Bioregulation (MIB), Kyushu University for the RNA-sequencing.

## Data Availability

RNA-seq data have been deposited in the Gene Expression Omnibus as accession number GSE263849.

## References

Agarwal, V. R., Ashanullah, C. I., Simpson, E. R., & Bulun, S. E. (1997). Alternatively spliced transcripts of the aromatase cytochrome P450 (CYP19) gene in adipose tissue of women. J Clin Endocrinol Metab, 82(1), 70–74. doi:10.1210/jcem.82.1.3655

Almeida, M., Iyer, S., Martin-Millan, M., Bartell, S. M., Han, L., Ambrogini, E., . . . Manolagas, S. C. (2013). Estrogen receptor-α signaling in osteoblast progenitors stimulates cortical bone accrual. J Clin Invest, 123(1), 394–404. doi:10.1172/jci65910

Alpern, D., Gardeux, V., Russeil, J., Mangeat, B., Meireles-Filho, A. C. A., Breysse, R., . . . Deplancke, B. (2019). BRB-seq: ultra-affordable high-throughput transcriptomics enabled by bulk RNA barcoding and sequencing. Genome Biol, 20(1), 71. doi:10.1186/s13059-019-1671-x

Batsis, J. A., & Villareal, D. T. (2018). Sarcopenic obesity in older adults: aetiology, epidemiology and treatment strategies. Nat Rev Endocrinol, 14(9), 513–537. doi:10.1038/s41574-018-0062-9

Becker, T., Lipscombe, L., Narod, S., Simmons, C., Anderson, G. M., & Rochon, P. A. (2012). Systematic review of bone health in older women treated with aromatase inhibitors for early-stage breast cancer. J Am Geriatr Soc, 60(9), 1761–1767. doi:10.1111/j.1532-5415.2012.04107.x

Bilezikian, J. P., Morishima, A., Bell, J., & Grumbach, M. M. (1998). Increased bone mass as a result of estrogen therapy in a man with aromatase deficiency. N Engl J Med, 339(9), 599–603. doi:10.1056/nejm199808273390905

Bouxsein, M. L., Boyd, S. K., Christiansen, B. A., Guldberg, R. E., Jepsen, K. J., & Müller, R. (2010). Guidelines for assessment of bone microstructure in rodents using micro-computed tomography. J Bone Miner Res, 25(7), 1468–1486. doi:10.1002/jbmr.141

Brown, K. A., Iyengar, N. M., Zhou, X. K., Gucalp, A., Subbaramaiah, K., Wang, H., . . . Dannenberg, A. J. (2017). Menopause Is a Determinant of Breast Aromatase Expression and Its Associations With BMI, Inflammation, and Systemic Markers. J Clin Endocrinol Metab, 102(5), 1692–1701. doi:10.1210/jc.2016-3606

Bulun, S. E., & Simpson, E. R. (1994). Competitive reverse transcription-polymerase chain reaction analysis indicates that levels of aromatase cytochrome P450 transcripts in adipose tissue of buttocks, thighs, and abdomen of women increase with advancing age. J Clin Endocrinol Metab, 78(2), 428–432. doi:10.1210/jcem.78.2.8106632

Carani, C., Qin, K., Simoni, M., Faustini-Fustini, M., Serpente, S., Boyd, J., . . . Simpson, E. R. (1997). Effect of testosterone and estradiol in a man with aromatase deficiency. N Engl J Med, 337(2), 91–95. doi:10.1056/nejm199707103370204

Dacquin, R., Starbuck, M., Schinke, T., & Karsenty, G. (2002). Mouse alpha1(I)-collagen promoter is the best known promoter to drive efficient Cre recombinase expression in osteoblast. Dev Dyn, 224(2), 245–251. doi:10.1002/dvdy.10100

Doolittle, M. L., Saul, D., Kaur, J., Rowsey, J. L., Eckhardt, B., Vos, S., . . . Khosla, S. (2022). Skeletal Effects of Inducible ERalpha Deletion in Osteocytes in Adult Mice. J Bone Miner Res, 37(9), 1750–1760. doi:10.1002/jbmr.4644

Dupé, V., Davenne, M., Brocard, J., Dollé, P., Mark, M., Dierich, A., . . . Rijli, F. M. (1997). In vivo functional analysis of the Hoxa-1 3’ retinoic acid response element (3’RARE). Development, 124(2), 399–410. doi:10.1242/dev.124.2.399

Dupont, S., Krust, A., Gansmuller, A., Dierich, A., Chambon, P., & Mark, M. (2000). Effect of single and compound knockouts of estrogen receptors alpha (ERalpha) and beta (ERbeta) on mouse reproductive phenotypes. Development, 127(19), 4277–4291.

Eicher, E. M., Southard, J. L., Scriver, C. R., & Glorieux, F. H. (1976). Hypophosphatemia: mouse model for human familial hypophosphatemic (vitamin D-resistant) rickets. Proc Natl Acad Sci U S A, 73(12), 4667–4671. doi:10.1073/pnas.73.12.4667

Emont, M. P., Jacobs, C., Essene, A. L., Pant, D., Tenen, D., Colleluori, G., . . . Rosen, E. D. (2022). A single-cell atlas of human and mouse white adipose tissue. Nature, 603(7903), 926–933. doi:10.1038/s41586-022-04518-2

Finkelstein, J. S., Lee, H., Burnett-Bowie, S. A., Pallais, J. C., Yu, E. W., Borges, L. F., . . . Leder, B. Z. (2013). Gonadal steroids and body composition, strength, and sexual function in men. N Engl J Med, 369(11), 1011–1022. doi:10.1056/NEJMoa1206168

Finkelstein, J. S., Lee, H., Leder, B. Z., Burnett-Bowie, S. A., Goldstein, D. W., Hahn, C. W., . . . Yu, E. W. (2016). Gonadal steroid-dependent effects on bone turnover and bone mineral density in men. J Clin Invest, 126(3), 1114–1125. doi:10.1172/JCI84137

Fisher, C. R., Graves, K. H., Parlow, A. F., & Simpson, E. R. (1998). Characterization of mice deficient in aromatase (ArKO) because of targeted disruption of the cyp19 gene. Proc Natl Acad Sci U S A, 95(12), 6965–6970. doi:10.1073/pnas.95.12.6965

Franco, L. P., Morais, C. C., & Cominetti, C. (2016). Normal-weight obesity syndrome: diagnosis, prevalence, and clinical implications. Nutr Rev, 74(9), 558–570. doi:10.1093/nutrit/nuw019

Fukumoto, S., & Martin, T. J. (2009). Bone as an endocrine organ. Trends Endocrinol Metab, 20(5), 230–236. doi:10.1016/j.tem.2009.02.001

Ghosh, D., Griswold, J., Erman, M., & Pangborn, W. (2009). Structural basis for androgen specificity and oestrogen synthesis in human aromatase. Nature, 457(7226), 219–223. doi:10.1038/nature07614

Goss, P. E., Ingle, J. N., Martino, S., Robert, N. J., Muss, H. B., Piccart, M. J., . . . Pater, J. L. (2003). A randomized trial of letrozole in postmenopausal women after five years of tamoxifen therapy for early-stage breast cancer. N Engl J Med, 349(19), 1793–1802. doi:10.1056/NEJMoa032312

Hanazaki, A., Ikuta, K., Sasaki, S., Sasaki, S., Koike, M., Tanifuji, K., . . . Segawa, H. (2020). Role of sodium-dependent Pi transporter/Npt2c on Pi homeostasis in klotho knockout mice different properties between juvenile and adult stages. Physiol Rep, 8(3), e14324. doi:10.14814/phy2.14324

Hayashibara, T., Hiraga, T., Sugita, A., Wang, L., Hata, K., Ooshima, T., & Yoneda, T. (2007). Regulation of osteoclast differentiation and function by phosphate: potential role of osteoclasts in the skeletal abnormalities in hypophosphatemic conditions. J Bone Miner Res, 22(11), 1743–1751. doi:10.1359/jbmr.070709

He, W., Barak, Y., Hevener, A., Olson, P., Liao, D., Le, J., . . . Evans, R. M. (2003). Adipose-specific peroxisome proliferator-activated receptor gamma knockout causes insulin resistance in fat and liver but not in muscle. Proc Natl Acad Sci U S A, 100(26), 15712–15717. doi:10.1073/pnas.2536828100

Honda, S., Harada, N., & Takagi, Y. (1994). Novel exon 1 of the aromatase gene specific for aromatase transcripts in human brain. Biochem Biophys Res Commun, 198(3), 1153–1160. doi:10.1006/bbrc.1994.1163

Huang da, W., Sherman, B. T., & Lempicki, R. A. (2009). Systematic and integrative analysis of large gene lists using DAVID bioinformatics resources. Nat Protoc, 4(1), 44–57. doi:10.1038/nprot.2008.211

Ikuta, K., Segawa, H., Hanazaki, A., Fujii, T., Kaneko, I., Shiozaki, Y., . . . Miyamoto, K. I. (2019). Systemic network for dietary inorganic phosphate adaptation among three organs. Pflugers Arch, 471(1), 123–136. doi:10.1007/s00424-018-2242-9

Ikuta, K., Segawa, H., Sasaki, S., Hanazaki, A., Fujii, T., Kushi, A., . . . Miyamoto, K. I. (2018). Effect of Npt2b deletion on intestinal and renal inorganic phosphate (Pi) handling. Clin Exp Nephrol, 22(3), 517–528. doi:10.1007/s10157-017-1497-3

Jones, M. E., Thorburn, A. W., Britt, K. L., Hewitt, K. N., Wreford, N. G., Proietto, J., . . . Simpson, E. R. (2000). Aromatase-deficient (ArKO) mice have a phenotype of increased adiposity. Proc Natl Acad Sci U S A, 97(23), 12735–12740. doi:10.1073/pnas.97.23.12735

Kaneko, I., Segawa, H., Ikuta, K., Hanazaki, A., Fujii, T., Tatsumi, S., . . . Miyamoto, K. I. (2018). Eldecalcitol Causes FGF23 Resistance for Pi Reabsorption and Improves Rachitic Bone Phenotypes in the Male Hyp Mouse. Endocrinology, 159(7), 2741–2758. doi:10.1210/en.2018-00109

Kim, N. R., David, K., Corbeels, K., Khalil, R., Antonio, L., Schollaert, D., . . . Dubois, V. (2021). Testosterone Reduces Body Fat in Male Mice by Stimulation of Physical Activity Via Extrahypothalamic ERalpha Signaling. Endocrinology, 162(6). doi:10.1210/endocr/bqab045

Kitajima, Y., & Ono, Y. (2016). Estrogens maintain skeletal muscle and satellite cell functions. J Endocrinol, 229(3), 267–275. doi:10.1530/joe-15-0476

Kondoh, S., Inoue, K., Igarashi, K., Sugizaki, H., Shirode-Fukuda, Y., Inoue, E., . . . Imai, Y. (2014). Estrogen receptor α in osteocytes regulates trabecular bone formation in female mice. Bone, 60, 68–77. doi:10.1016/j.bone.2013.12.005

Lu, Y., Xie, Y., Zhang, S., Dusevich, V., Bonewald, L. F., & Feng, J. Q. (2007). DMP1-targeted Cre expression in odontoblasts and osteocytes. J Dent Res, 86(4), 320–325. doi:10.1177/154405910708600404

Lunenfeld, B., Saad, F., & Hoesl, C. E. (2005). ISA, ISSAM and EAU recommendations for the investigation, treatment and monitoring of late-onset hypogonadism in males: scientific background and rationale. Aging Male, 8(2), 59–74. doi:10.1080/13685530500163416

Mahendroo, M. S., Mendelson, C. R., & Simpson, E. R. (1993). Tissue-specific and hormonally controlled alternative promoters regulate aromatase cytochrome P450 gene expression in human adipose tissue. J Biol Chem, 268(26), 19463–19470.

Mair, K. M., Gaw, R., & MacLean, M. R. (2020). Obesity, estrogens and adipose tissue dysfunction - implications for pulmonary arterial hypertension. Pulm Circ, 10(3), 2045894020952019. doi:10.1177/2045894020952023

Martin-Millan, M., Almeida, M., Ambrogini, E., Han, L., Zhao, H., Weinstein, R. S., . . . Manolagas, S. C. (2010). The estrogen receptor-alpha in osteoclasts mediates the protective effects of estrogens on cancellous but not cortical bone. Mol Endocrinol, 24(2), 323–334. doi:10.1210/me.2009-0354

Means, G. D., Kilgore, M. W., Mahendroo, M. S., Mendelson, C. R., & Simpson, E. R. (1991). Tissue-specific promoters regulate aromatase cytochrome P450 gene expression in human ovary and fetal tissues. Mol Endocrinol, 5(12), 2005–2013. doi:10.1210/mend-5-12-2005

Mellström, D., Vandenput, L., Mallmin, H., Holmberg, A. H., Lorentzon, M., Odén, A., . . . Ohlsson, C. (2008). Older men with low serum estradiol and high serum SHBG have an increased risk of fractures. J Bone Miner Res, 23(10), 1552–1560. doi:10.1359/jbmr.080518

Minisola, S., Peacock, M., Fukumoto, S., Cipriani, C., Pepe, J., Tella, S. H., & Collins, M. T. (2017). Tumour-induced osteomalacia. Nat Rev Dis Primers, 3, 17044. doi:10.1038/nrdp.2017.44

Miyaura, C., Toda, K., Inada, M., Ohshiba, T., Matsumoto, C., Okada, T., . . . Ito, A. (2001). Sex- and age-related response to aromatase deficiency in bone. Biochem Biophys Res Commun, 280(4), 1062–1068. doi:10.1006/bbrc.2001.4246

Morishima, A., Grumbach, M. M., Simpson, E. R., Fisher, C., & Qin, K. (1995). Aromatase deficiency in male and female siblings caused by a novel mutation and the physiological role of estrogens. J Clin Endocrinol Metab, 80(12), 3689–3698. doi:10.1210/jcem.80.12.8530621

Nakamura, T., Imai, Y., Matsumoto, T., Sato, S., Takeuchi, K., Igarashi, K., . . . Kato, S. (2007). Estrogen prevents bone loss via estrogen receptor alpha and induction of Fas ligand in osteoclasts. Cell, 130(5), 811–823. doi:10.1016/j.cell.2007.07.025

Ohlsson, C., Hammarstedt, A., Vandenput, L., Saarinen, N., Ryberg, H., Windahl, S. H., . . . Sjogren, K. (2017). Increased adipose tissue aromatase activity improves insulin sensitivity and reduces adipose tissue inflammation in male mice. Am J Physiol Endocrinol Metab, 313(4), E450–E462. doi:10.1152/ajpendo.00093.2017

Oz, O. K., Hirasawa, G., Lawson, J., Nanu, L., Constantinescu, A., Antich, P. P., . . . Simpson, E. R. (2001). Bone phenotype of the aromatase deficient mouse. J Steroid Biochem Mol Biol, 79(1-5), 49–59. doi:10.1016/s0960-0760(01)00130-3

Pacifici, R. (2008). Estrogen deficiency, T cells and bone loss. Cell Immunol, 252(1-2), 68–80. doi:10.1016/j.cellimm.2007.06.008

Picelli, S., Björklund, A. K., Reinius, B., Sagasser, S., Winberg, G., & Sandberg, R. (2014). Tn5 transposase and tagmentation procedures for massively scaled sequencing projects. Genome Res, 24(12), 2033–2040. doi:10.1101/gr.177881.114

Prieto, C., & Barrios, D. (2020). RaNA-Seq: Interactive RNA-Seq analysis from FASTQ files to functional analysis. Bioinformatics, 36(6), 1955–1956. doi:10.1093/bioinformatics/btz854

Rochira, V., & Carani, C. (2009). Aromatase deficiency in men: a clinical perspective. Nat Rev Endocrinol, 5(10), 559–568. doi:10.1038/nrendo.2009.176

Rodda, S. J., & McMahon, A. P. (2006). Distinct roles for Hedgehog and canonical Wnt signaling in specification, differentiation and maintenance of osteoblast progenitors. Development, 133(16), 3231–3244. doi:10.1242/dev.02480

Rosen, C. J. (2005). Clinical practice. Postmenopausal osteoporosis. N Engl J Med, 353(6), 595–603. doi:10.1056/NEJMcp043801

Sakai, H., Sawada, Y., Tokunaga, N., Tanaka, K., Nakagawa, S., Sakakibara, I., Imai, Y. (2022). Uhrf1 governs the proliferation and differentiation of muscle satellite cells. iScience, 25(3), 103928. doi:10.1016/j.isci.2022.103928

Sakai, H., Uno, H., Yamakawa, H., Tanaka, K., Ikedo, A., Uezumi, A., . . . Imai, Y. (2024). The androgen receptor in mesenchymal progenitors regulates skeletal muscle mass via Igf1 expression in male mice. Proc Natl Acad Sci U S A, 121(39), e2407768121. doi:10.1073/pnas.2407768121

Sakakibara, I., Yanagihara, Y., Himori, K., Yamada, T., Sakai, H., Sawada, Y., . . . Imai, Y. (2021). Myofiber androgen receptor increases muscle strength mediated by a skeletal muscle splicing variant of Mylk4. iScience, 24(4), 102303. doi:10.1016/j.isci.2021.102303

Sasano, H., Uzuki, M., Sawai, T., Nagura, H., Matsunaga, G., Kashimoto, O., & Harada, N. (1997). Aromatase in human bone tissue. J Bone Miner Res, 12(9), 1416–1423. doi:10.1359/jbmr.1997.12.9.1416

Sato, S., Arimura, Y., Kujirai, T., Harada, A., Maehara, K., Nogami, J., . . . Kurumizaka, H. (2019). Biochemical analysis of nucleosome targeting by Tn5 transposase. Open Biol, 9(8), 190116. doi:10.1098/rsob.190116

Scheller, E. L., Troiano, N., Vanhoutan, J. N., Bouxsein, M. A., Fretz, J. A., Xi, Y., . . . Horowitz, M. C. (2014). Use of osmium tetroxide staining with microcomputerized tomography to visualize and quantify bone marrow adipose tissue in vivo. Methods Enzymol, 537, 123–139. doi:10.1016/B978-0-12-411619-1.00007-0

Segawa, H., Onitsuka, A., Furutani, J., Kaneko, I., Aranami, F., Matsumoto, N., . . . Miyamoto, K. (2009). Npt2a and Npt2c in mice play distinct and synergistic roles in inorganic phosphate metabolism and skeletal development. Am J Physiol Renal Physiol, 297(3), F671–678. doi:10.1152/ajprenal.00156.2009

Segawa, H., Yamanaka, S., Ito, M., Kuwahata, M., Shono, M., Yamamoto, T., & Miyamoto, K. (2005). Internalization of renal type IIc Na-Pi cotransporter in response to a high-phosphate diet. Am J Physiol Renal Physiol, 288(3), F587–596. doi:10.1152/ajprenal.00097.2004

Segawa, H., Yamanaka, S., Onitsuka, A., Tomoe, Y., Kuwahata, M., Ito, M., . . . Miyamoto, K. (2007). Parathyroid hormone-dependent endocytosis of renal type IIc Na-Pi cotransporter. Am J Physiol Renal Physiol, 292(1), F395–403. doi:10.1152/ajprenal.00100.2006

Shahinian, V. B., Kuo, Y. F., Freeman, J. L., & Goodwin, J. S. (2005). Risk of fracture after androgen deprivation for prostate cancer. N Engl J Med, 352(2), 154–164. doi:10.1056/NEJMoa041943

Sherman, B. T., Hao, M., Qiu, J., Jiao, X., Baseler, M. W., Lane, H. C., . . . Chang, W. (2022). DAVID: a web server for functional enrichment analysis and functional annotation of gene lists (2021 update). Nucleic Acids Res, 50(W1), W216–w221. doi:10.1093/nar/gkac194

Shimada, T., Hasegawa, H., Yamazaki, Y., Muto, T., Hino, R., Takeuchi, Y., . . . Yamashita, T. (2004). FGF-23 is a potent regulator of vitamin D metabolism and phosphate homeostasis. J Bone Miner Res, 19(3), 429–435. doi:10.1359/jbmr.0301264

Shozu, M., & Simpson, E. R. (1998). Aromatase expression of human osteoblast-like cells. Mol Cell Endocrinol, 139(1-2), 117–129. doi:10.1016/s0303-7207(98)00069-0

Simpson, E. R., Mahendroo, M. S., Means, G. D., Kilgore, M. W., Corbin, C. J., & Mendelson, C. R. (1993). Tissue-specific promoters regulate aromatase cytochrome P450 expression. J Steroid Biochem Mol Biol, 44(4-6), 321–330. doi:10.1016/0960-0760(93)90235-o

Sun, L., Peng, Y., Sharrow, A. C., Iqbal, J., Zhang, Z., Papachristou, D. J., . . . Zaidi, M. (2006). FSH directly regulates bone mass. Cell, 125(2), 247–260. doi:10.1016/j.cell.2006.01.051

Takashi, Y. (2024). Phosphate-sensing mechanisms and functions of phosphate as a first messenger. Endocr J. doi:10.1507/endocrj.EJ24-0082

Takashi, Y., Kosako, H., Sawatsubashi, S., Kinoshita, Y., Ito, N., Tsoumpra, M. K., . . . Fukumoto, S. (2019). Activation of unliganded FGF receptor by extracellular phosphate potentiates proteolytic protection of FGF23 by its O-glycosylation. Proc Natl Acad Sci U S A, 116(23), 11418–11427. doi:10.1073/pnas.1815166116

Takashi, Y., Sawatsubashi, S., Endo, I., Ohnishi, Y., Abe, M., Matsuhisa, M., . . . Fukumoto, S. (2021). Skeletal FGFR1 signaling is necessary for regulation of serum phosphate level by FGF23 and normal life span. Biochem Biophys Rep, 27, 101107. doi:10.1016/j.bbrep.2021.101107

Walton, R. J., & Bijvoet, O. L. (1975). Nomogram for derivation of renal threshold phosphate concentration. Lancet, 2(7929), 309–310. doi:10.1016/s0140-6736(75)92736-1

Yanase, T., Suzuki, S., Goto, K., Nomura, M., Okabe, T., Takayanagi, R., & Nawata, H. (2003). Aromatase in bone: roles of Vitamin D3 and androgens. The Journal of Steroid Biochemistry and Molecular Biology, 86(3-5), 393–397. doi:10.1016/s0960-0760(03)00349-2

Zhang, M., Xuan, S., Bouxsein, M. L., von Stechow, D., Akeno, N., Faugere, M. C., . . . Clemens, T. L. (2002). Osteoblast-specific knockout of the insulin-like growth factor (IGF) receptor gene reveals an essential role of IGF signaling in bone matrix mineralization. J Biol Chem, 277(46), 44005–44012. doi:10.1074/jbc.M208265200

