## Supplemental information for "Aromatase in adipose tissue exerts an osteoprotective function in male mice via phosphate regulation"

Figure S1

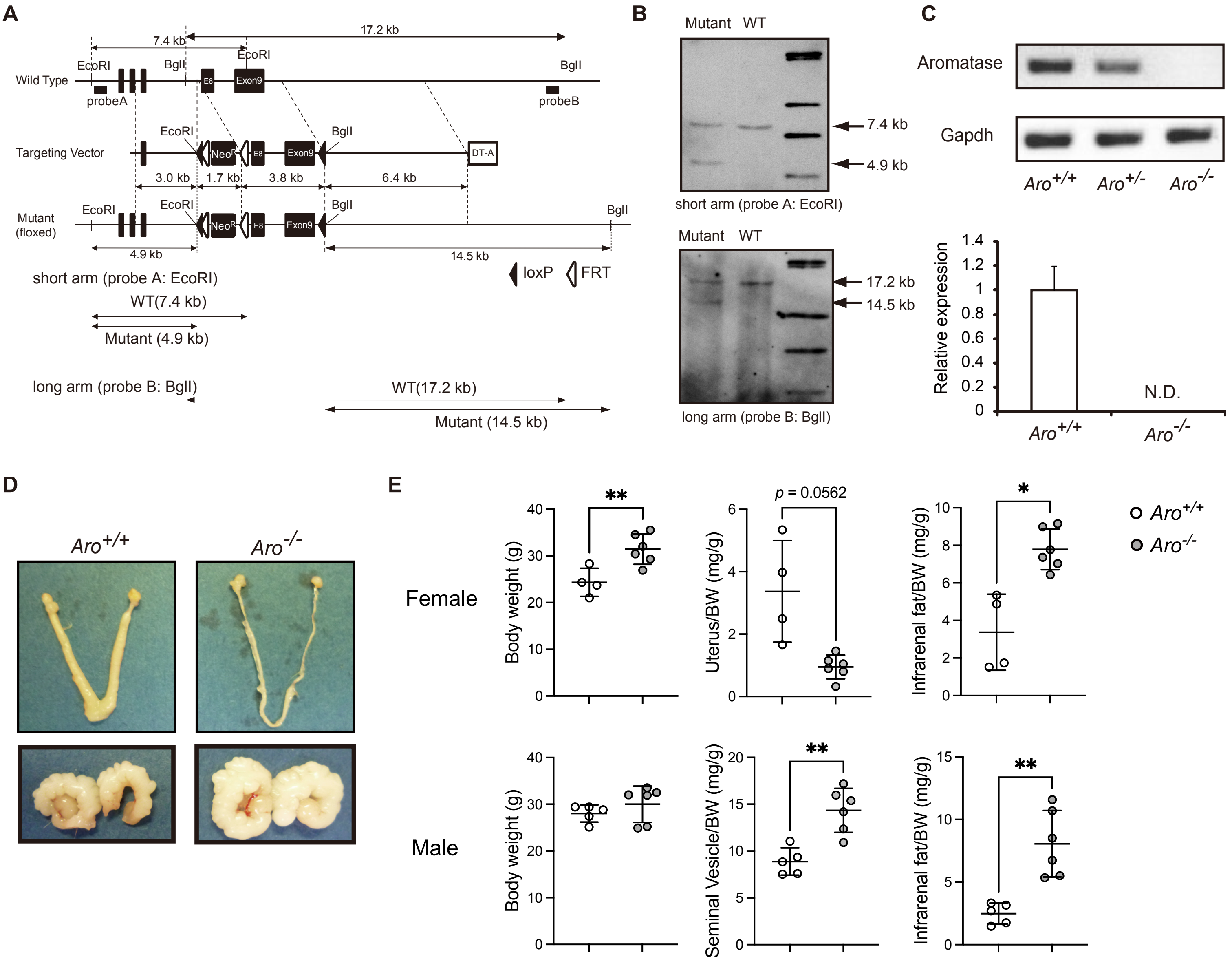


**Figure S1.** Generation of *Cyp19a1* floxed mice and systemic *Cyp19a1* KO mice (*Aro*$^{-/-}$). (A) Targeting strategy with positive-/negative- selection. The strategy of genomic Southern blotting in the screening for homologous recombinant embryonic stem (ES) cell clones is also included. E8 represents exon 8 of the *Cyp19a1* gene. The solid triangles represent the LoxP sites. The open triangles represent Frt sites. (B) Southern blotting analysis of targeted ES clones. EcoRI was used to screen recombination events with probe A for the short arm. A 7.4 kb fragment in the wild-type and a 4.9 kb fragment after homologous recombination were confirmed with probe A (upper panel). BglI was used for screening recombination events with probe B. A 17.2 kb fragment in the wild-type and a 14.5 kb fragment after homologous recombination were confirmed with probe B (lower panel). (C) Qualitative (upper panels) and quantitative (lower panel) RT-PCR were performed to detect the transcript of the Cyp19a1 gene, aromatase (*Aro*), using total RNA extracted from the brains of control (*Aro*$^{+/+}$) mice, and homozygous Ar knockout (*Aro*$^{-/-}$) mice with or without heterozygous Aro deficient (*Aro*$^{+/-}$) littermates. Data are presented as means ± SEM. (D) Representative views of uteri (upper panels) and seminal vesicles (lower panels) of control and *Aro*$^{-/-}$ mice. (E) Weights of uteri, infrarenal fat and seminal vesicles of control ($n$ = 4 or 5) and *Aro*$^{-/-}$ mice ($n$ = 6). Data are presented as means ± SD except in S1 C. *, $p < 0.05$; **, $p < 0.01$. N.D.; not detected.

Figure S2

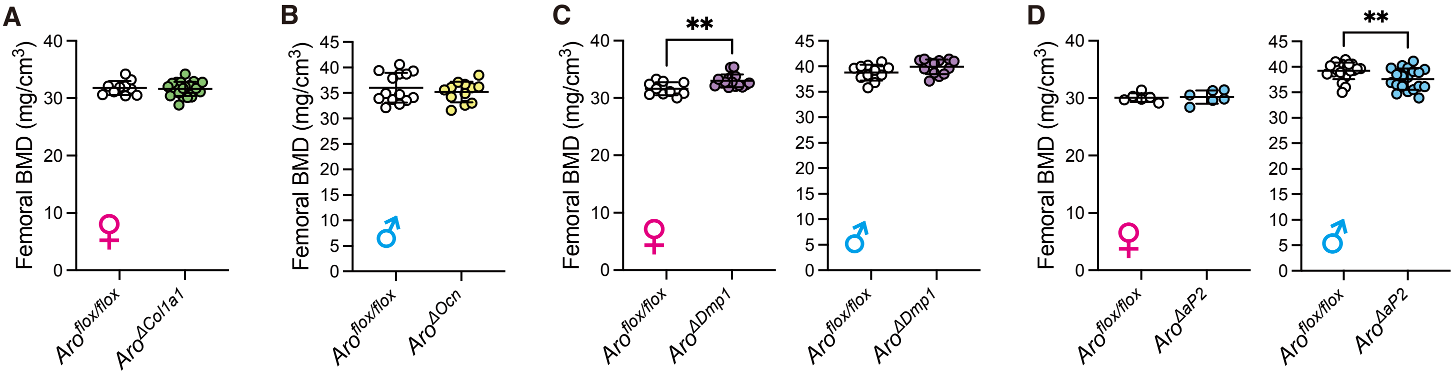


**Figure S2.** Three lines of osteoblast/osteocyte-specific *Cyp19a1* knockout mice were created. Femoral BMD in (A) *Aro*$^{\Delta Col1a1}$, (B) *Aro*$^{\Delta OCN}$, (C) *Aro*$^{\Delta Dmp1}$, and (D) *Aro*$^{\Delta aP2}$ mice and control mice for each are shown. Data are presented as means ± SD. **, $p < 0.01$.

Figure S3

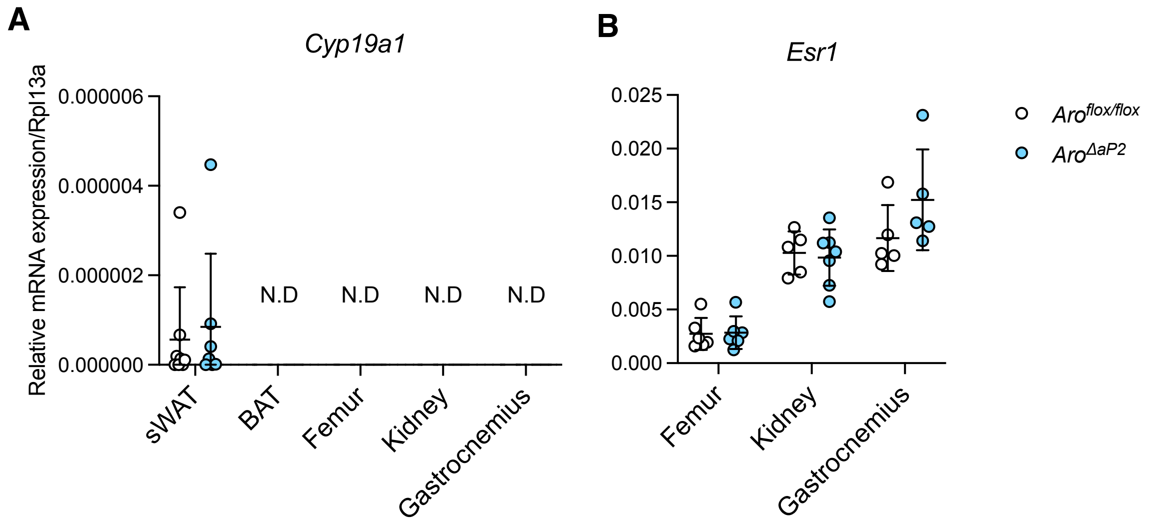


**Figure S3.** Expression levels of *Cyp19a1* and *Esr1* in sWAT, BAT, bone, kidney, and muscle. (A) *Cyp19a1* expressin levels in sWAT, BAT, femoral bone, kidney, and gastrocnemius muscle. (B) *Esr1* expressin levels in femoral bone, kidney, and gastrocnemius muscle. Data are presented as means ± SD.

Figure S4

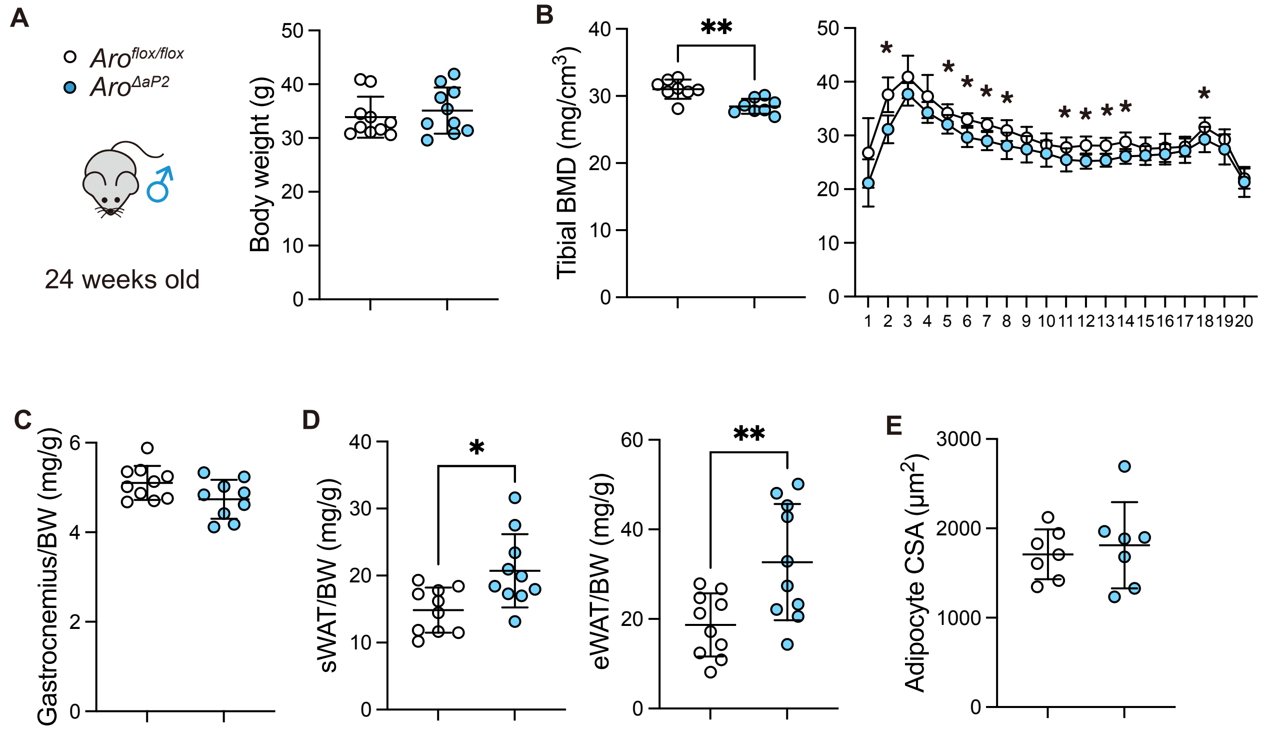


**Figure S4.** Male *Aro*$^{\Delta aP2}$ mice at 24 weeks old was also exhibited low bone mass and high fat mass. (A) Body weight (BW). (B) Tibial bone mineral density (BMD). (C) Gastrocnemius muscle weight per BW. (D) Subcutaneous and epidydimal white adipose tissue (WAT) weight per BW. (E) Adypocyte CSA of eWAT. Data are presented as means ± SD.

Figure S5

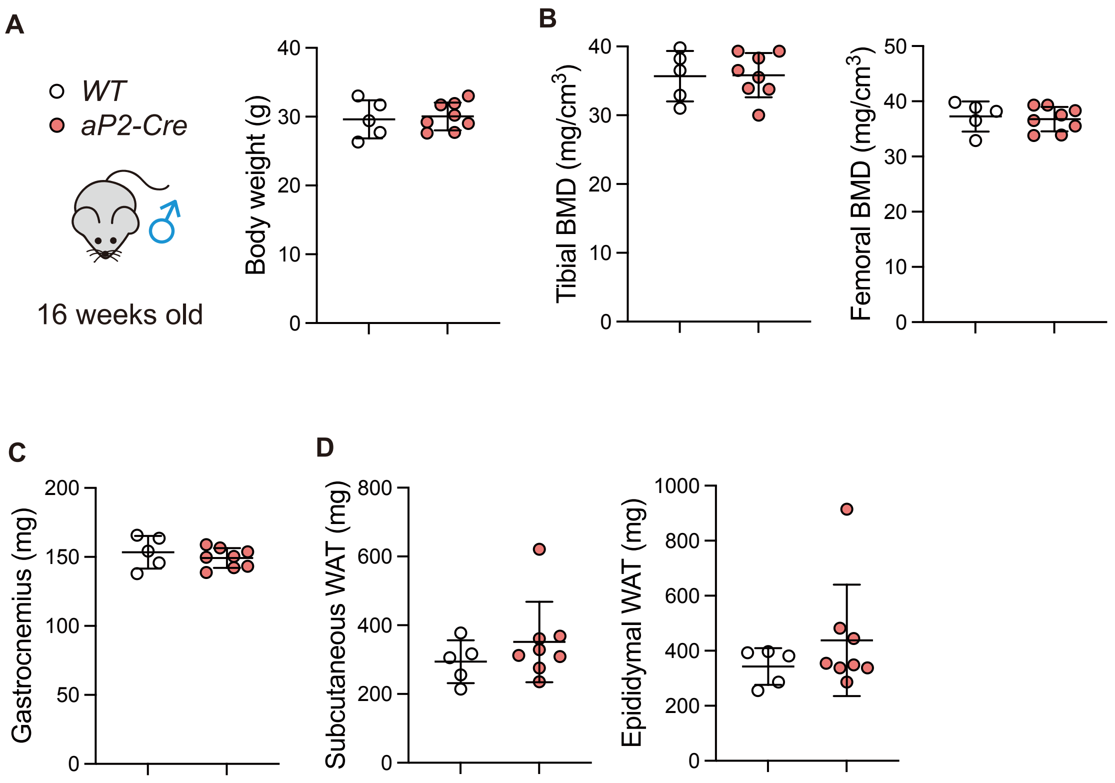


**Figure S5.** Confirmation of *aP2-Cre* mouse phenotype. (A) Body weight. (B) Tibial and femoral bone mineral density (BMD). (C) Gastrocnemius muscle weight. (D) Subcutaneous and epidydimal white adipose tissue (WAT) weight. Data are presented as means ± SD.

Fugure S6

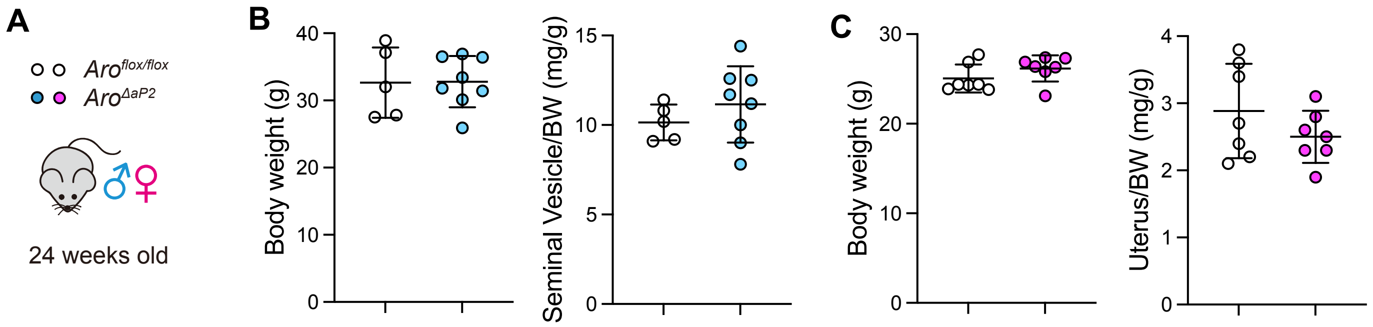


**Figure S6.** Seminal vesicle weight in male and uterus weight in female. (A) Twenty-four week old *Aro*$^{ΔaP2}$ and *Aro*$^{flox/flox}$ mice were analyzed. (B) Body weight and seminal vesicle weight in male. (C) Body weight and uterus weight in female. Data are presented as means ± SD.

Figure S7

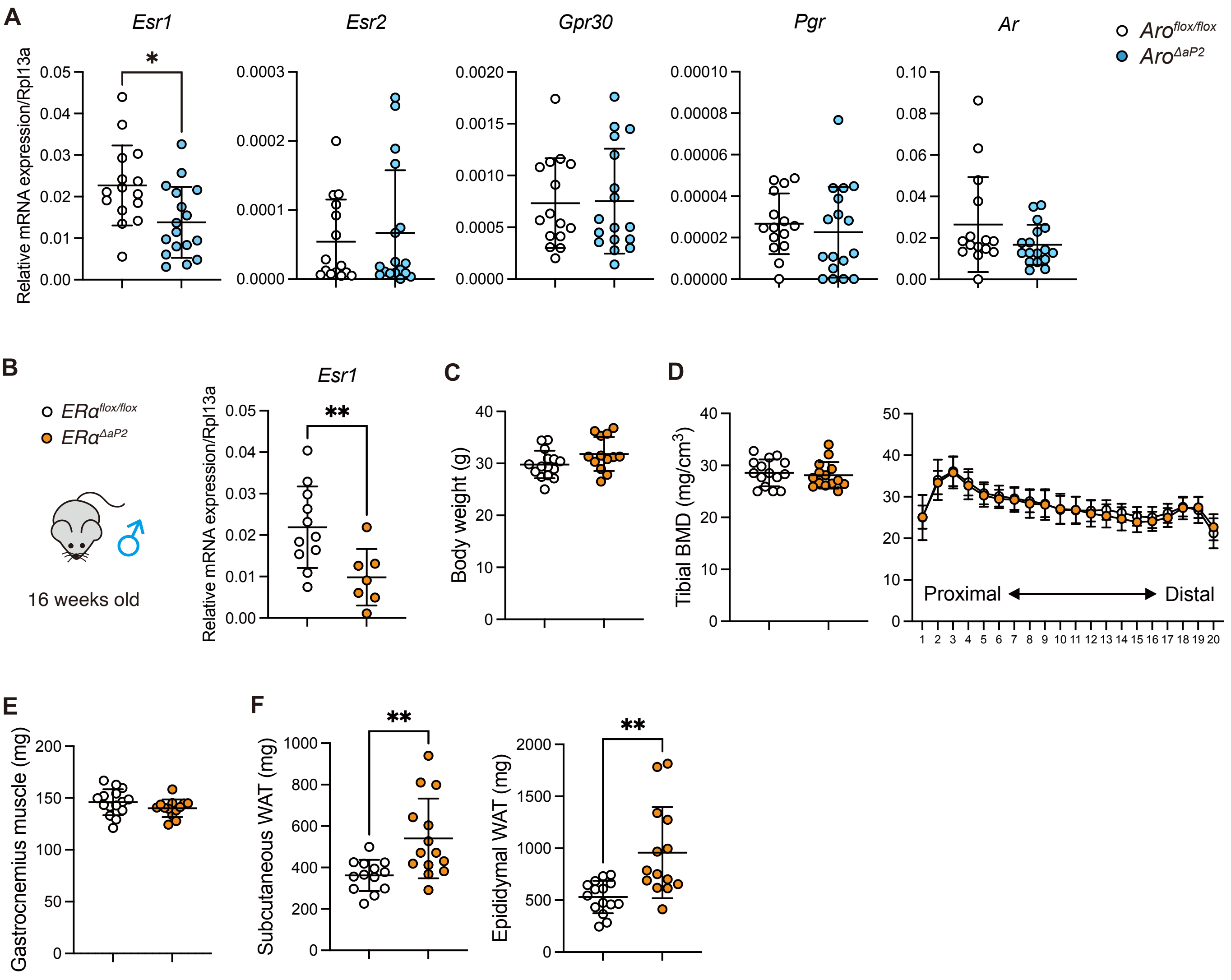


**Figure S7.** Steroid hormone receptor expression in eWAT of $Aro^{\Delta aP2}$ mice and phenotype of adipocyte-specific ER α KO mice ($ER\alpha^{\Delta aP2}$). (A) Gene expression of *Esr1*, *Esr2*, *Gpr30*, *Pgr*, and *Ar* in eWAT. (B) *Esr1* expression level in eWAT of $ER\alpha^{\Delta aP2}$. (C) Body weight. (D) Total tibial BMD (left) and BMD at each site when the bone is divided into 20 sections (right). (E) Gastrocnemius muscle weight. (F) Subcutaneous and epidydimal white adipose tissue (WAT) weight. Data are presented as means ± SD. *, $p < 0.05$; **, $p < 0.01$

Figure S8

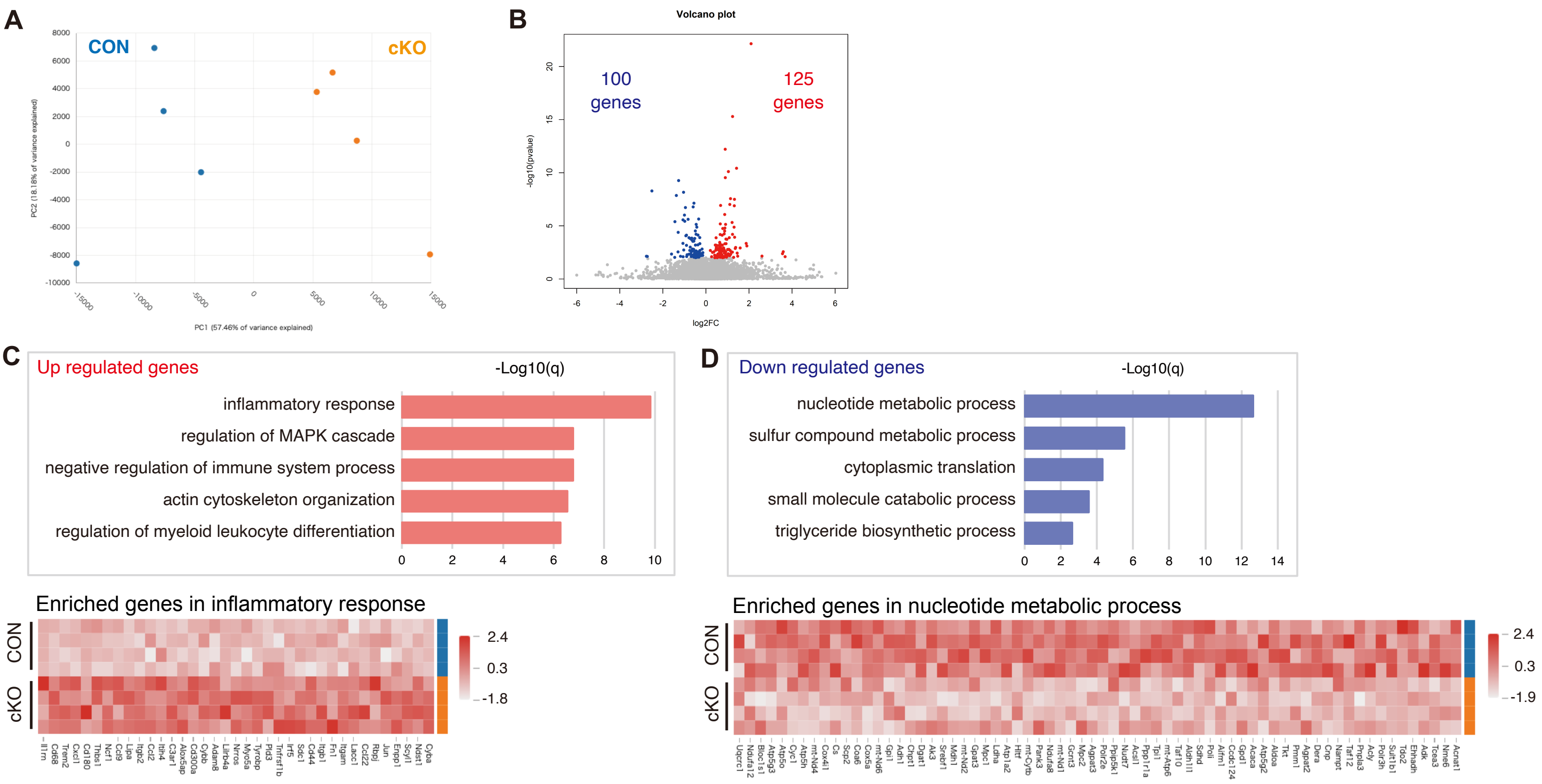


**Figure S8.** RNA-seq of eWAT in 16 weeks old *Aro*$^{\Delta aP2}$ and *Aro*$^{flox/flox}$ mice. (A) Principal Components Analysis (PCA). (B) Volcano plot ($p < 0.01$). (C) Results of GO analysis of up regulated genes and (D) down regulated genes.

Figure S9

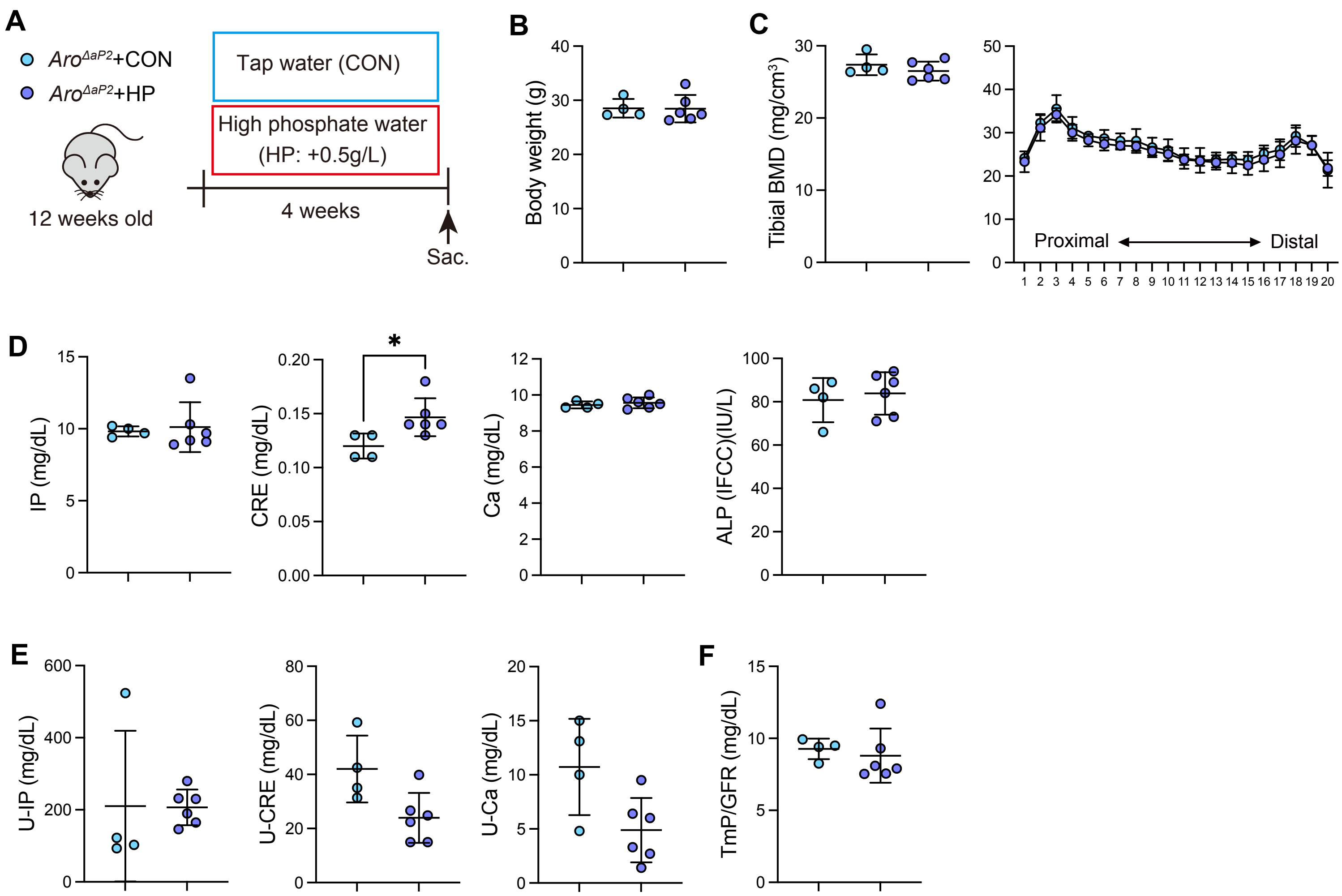


**Figure S9.** High phosphate warter expreiment. (A) $Aro^{\Delta aP2}$ and $Aro^{flox/flox}$ mice at 12 weeks old were fed with high-phosphate water (HP) or tap water (CON) for 4 weeks. (B) Body weight. (C) Total tibial BMD (left) and BMD at each site when the bone is divided into 20 sections (right). (D) Serum IP, CRE, Ca, and ALP. (E) Urine IP, CRE, and Ca. (F) Tmp/GFR calculated by serum and urine parameters. Data are presented as means ± SD. *, $p < 0.05$; **, $p < 0.01$

**Table S1.** qPCR primers

| Target | Primer sequence 5'-> 3' | |
|---|---|---|
| | Forward | Reverse |
| *Gapdh* | TGTGTCCGTCGTGGATCTGA | TTGCTGTGAAGTCGCAGGAG |
| *Rpl13a* | GTGGTCCCTGCTGCTCTCAAG | CGATAGTGCATCTTGGCCTTTT |
| *Cyp19a1* | GGTCGAAGCAGCAATCCTGAA | AGAGCTCTGCGCATGACCAA |
| *Npt2a* | AGTCTCATTCGGATTTGGTGTCA | GCCGATGGCCTCTACCCT |
| *Npt2c* | TAATCTTCGCAGTTCAGGTTGCT | CAGTGGAATTGGCAGTCTCAA |
| *Esr1* | CCGCCTTCTACAGGTCTAATTC | AGGCATAGTCATTGCACACG |
| *Esr2* | CTGTGATGAACTACAGTGTTCCC | CACATTTGGGCTTGCAGTCTG |
| *Gpr30* | GCCTCTGCTACTCCCTCATC | ACTGCGAAGATCATCCTCAGG |
| *Pgr* | CTCCGGGACCGAACAGAGT | ACAACAACCCTTTGGTAGCAG |
